# Hepatic WDR23 proteostasis mediates insulin clearance by regulating insulin-degrading enzyme

**DOI:** 10.1101/2022.11.24.516014

**Authors:** Chatrawee Duangjan, Thalida Em Arpawong, Brett N. Spatola, Sean P. Curran

**Affiliations:** Leonard Davis School of Gerontology, University of Southern California, Los Angeles, CA 90089; 2Dornsife College of Letters, Arts, and Science, University of Southern California, Los Angeles, CA 90089

**Keywords:** Insulin-degrading enzyme (IDE), insulin clearance, WDR23, proteostasis, liver, hepatocytes, NRF2

## Abstract

Clearance of circulating insulin is critical for metabolic homeostasis. In the liver, insulin is degraded by the activity of the insulin-degrading enzyme (IDE). Here we establish a hepatic regulatory axis for IDE through WDR23-proteostasis. *Wdr23KO* mice have increased IDE expression, reduced circulating insulin, and defective insulin responses. Genetically engineered human cell models lacking *WDR23* also increase IDE expression and display dysregulated phosphorylation of insulin signaling cascade proteins, IRS-1, AKT2, MAPK, FoxO, and mTOR, similar to cells treated with insulin, which can be mitigated by chemical inhibition of IDE. Mechanistically, the cytoprotective transcription factor NRF2, a direct target of WDR23-Cul4 proteostasis, mediates the enhanced transcriptional expression of IDE when *WDR23* is ablated. Moreover, an analysis of human genetic variation in *WDR23* across a large naturally aging human cohort in the US Health and Retirement Study reveals a significant association of *WDR23* with altered hemoglobin A1C (HbA1c) levels in older adults, supporting the use of *WDR23* as new molecular determinant of metabolic health in humans.

## INTRODUCTION

The incidence of diabetes continues to increase with over one million new diagnoses each year. Unlike cases of type I diabetes where the individual does not produce an adequate amount of insulin, individuals with type 2 diabetes (T2D) do not effectively respond to the insulin produced. More than 90% of diabetes cases are type 2 and strikingly, 90 million new cases of pre-diabetes are documented each year [1, 2]. However, it is estimated that more than 20% of individuals with diabetes are unaware of their condition. Diabetes is diagnosed phenotypically in the clinic, and our understanding of the etiology of T2D and ability to predict a genetic predisposition, is currently limited by our understanding of the entirety of molecular regulators that influence this critical metabolic homeostat.

Following its secretion from Beta cells in the pancreas, clearance of endogenously released insulin is primarily achieved by hepatocytes in the liver [3]. ∼80% of released insulin is cleared in the first pass through the liver [4] while subsequent passage through the hepatic artery can further deplete insulin from circulation [5]. Defective insulin clearance has been linked to T2D [6] as well as hyperinsulinemia-driven systemic insulin resistance [7, 8] and hyperinsulinemia in metabolic syndrome [9]. In the obese state, hyperinsulinemia results from increased insulin secretion, but also from impaired clearance [10–12].

Insulin-degrading enzyme (IDE) is a ubiquitously expressed metalloprotease with a high affinity for insulin [6, 13]. IDE can degrade insulin in multiple intracellular compartments [14] and genetic ablation of *Ide* results in hyperinsulinemia which suggests IDE plays a central role in insulin clearance. However, non-proteolytic roles for IDE in insulin metabolism by downregulation of the insulin receptor have also been documented [15]. Despite these established roles in insulin metabolism, the regulatory mechanisms that govern IDE expression and activity are not fully understood.

The ubiquitin-proteasome system (UPS) is the primary protein degradation pathway, which plays an important role in cellular proteostasis [16, 17]. Proteins are targeted to the proteasome by a collection of ubiquitin-conjugating enzyme complexes, and poly-ubiquitinated target proteins are degraded by the proteasome [16]. Cullin-RING ligases (CRLs) are a well-known class of E3-ubiquitin ligases found in eukaryotes [17, 18], in which substrate receptors – including DDB1-CUL4 associated factors (DCAFs), also known as WD repeat (WDR) proteins - provide target specificity to the complex. However, the specific substrates for each receptor protein, and their functions in human health and disease, are still largely unknown. Previously, we defined WDR23 as the substrate receptor for the cytoprotective transcription factor NRF2 that functions independently to the canonical KEAP1-CUL3 regulatory pathway [19]. Moreover, GEN1 [17] and SLBP [20] are confirmed substrates of the WDR23-CUL4 proteostat, but additional substrates remain to be identified and are likely to play critical biological functions.

In the present study, we utilize a new *Wdr23KO* mouse model to expand upon our previous investigation of the physiological roles of WDR23, first studied in *C. elegans* [17, 19], and define a role for the WDR23 proteostasis pathway in insulin clearance and organismal metabolic homeostasis. We further define human genetic variation in WDR23 as a factor associated with diabetes. Taken together, our work defines WDR23 as a new factor in cellular and organismal insulin homeostasis.

## RESULTS

### Loss of WDR23 disrupts insulin sensitivity in male mice

To define the role of WDR23 proteostasis in vertebrate physiology, we commissioned the generation of a floxed allele of *Wdr23* to generate animals lacking *Wdr23* expression in all tissues; hereafter referred to as *Wdr23KO* (**Figure 1A**). *Wdr23KO* mice are viable, display no overt defects in sexual maturity or reproductive capacity, and display normal body weight in both sexes when compared to wild-type (WT) animals over 44 weeks on a standardized 10% fat diet (**Figure 1B-C** and Figure S1A-B).

**Figure 1.**
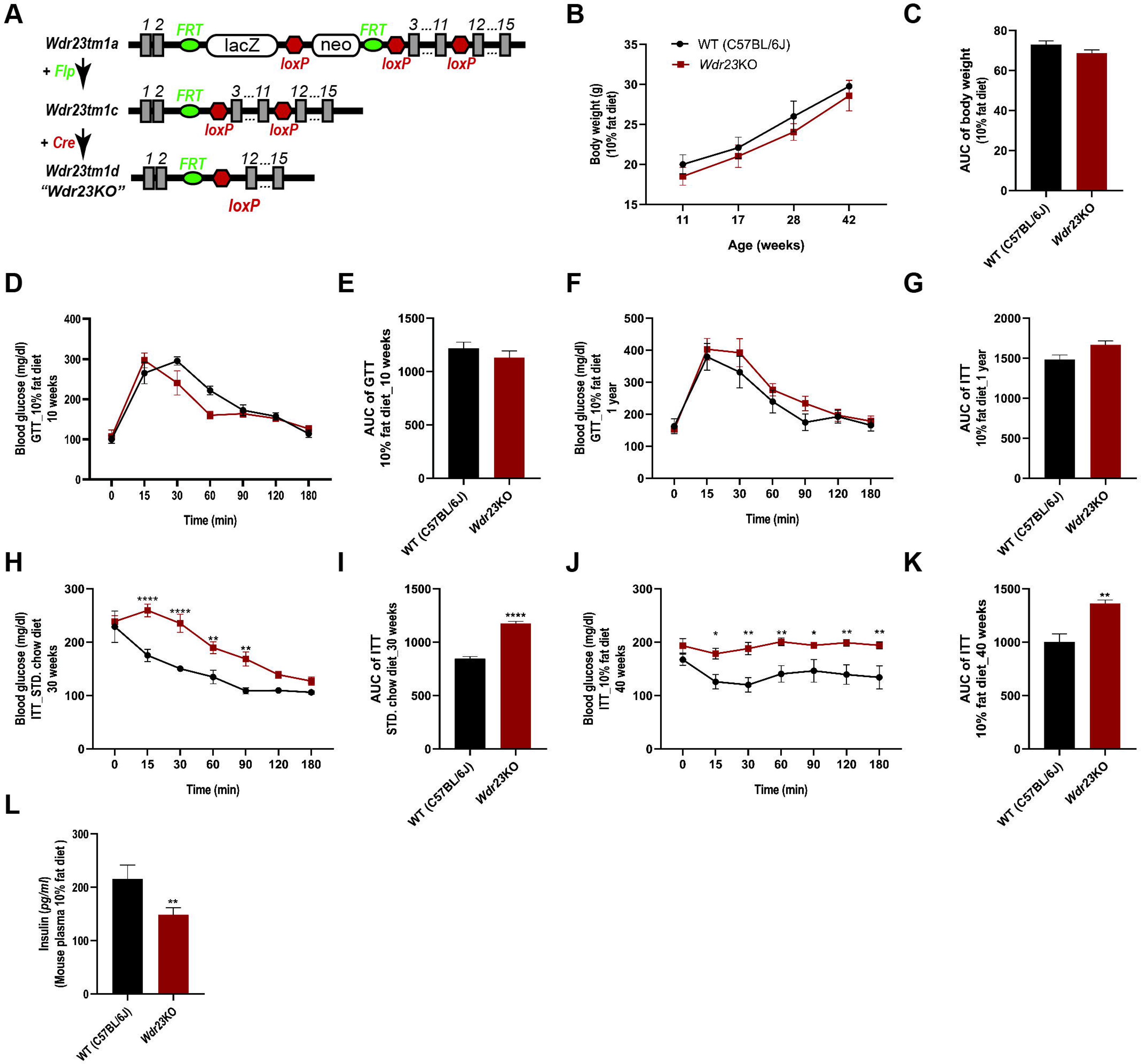
*Wdr23KO* male mice display impaired insulin homeostasis. Model of Cre-mediated germline deletion of *Wdr23* (**A**). *Wdr23KO* animals gain weight at similar rates as age-matched WT (C57BL/6J) animals (**B,C**). Glucose clearance, as measured by glucose tolerance testing (GTT) is similar between WT and *Wdr23KO* male mice at 10-weeks (**D,E**) and 1-year of age (**F,G**). Insulin tolerance is impaired in *Wdr23KO* male mice fed a standard chow diet (**H, I**) as well as animals fed a chemically defined 10% fat diet (**J, K**). ELISA analysis of mouse plasma quantifying circulating insulin levels (**L**). *p<.05, **p<.01, ***p<.001, ****p<.0001

We next examined the effect of *Wdr23* deletion on glucose homeostasis and insulin sensitivity by glucose tolerance test (GTT) and insulin tolerance test (ITT), respectively. At all ages tested, male *Wdr23KO* mice display an impairment of insulin sensitivity while exhibiting normal glucose clearance when compared to age-matched WT controls (**Figure 1D-K**). However, the effects of *Wdr23KO* were sexually dimorphic as neither glucose tolerance nor insulin sensitivity were different in female *Wdr23KO* mice as compared to WT (Figure S1C-J). As such, we used male mice in all subsequent experiments.

In light of the potential differences in the responsiveness toward ectopically delivered insulin and the use of endogenously produced insulin [21], we next examined whether steady-state insulin levels were impacted by the loss of *Wdr23*. Surprisingly, we noted a significant reduction in the levels of circulating insulin in *Wdr23KO* mice (**Figure 1L**).

Hepatic steatosis is associated with insulin resistance [21, 22] and as such, we next examined histological comparisons between the livers from 44-week-old mice. We observed no significant changes in liver morphology of *Wdr23KO* mice, compared to the WT mice (Figure S1K-P). Taken together, these data indicated that the loss of *Wdr23* results in early defects in insulin sensitivity; before glucose handling impairment and gross morphological defects in liver organization.

### *Wdr23*KO mice accumulate insulin-degrading enzyme (IDE) in the liver

In the ubiquitin proteasome system, loss of a substrate receptor leads to a loss in turnover of substrates associated with that receptor [17, 19, 23, 24]. To identify new substrates of WDR23 we performed an unbiased proteomic analysis of liver from WT and *Wdr23KO* mice. We examined the liver as this tissue plays a crucial role in the regulation of glucose homeostasis [21] and insulin sensitivity [25] and provided adequate sample mass for analysis. 209 unique proteins were identified with increased abundance across the samples; nine (9) proteins were classified as high confidence (>30% change, p<0.01) and 200 proteins were classified with moderate confidence (5-30% change, 0.01<p<0.05) (**Figure 2A**; Table S3). Among the nine high confidence proteins, we found that the level of insulin-degrading enzyme (IDE) was significantly increased in *Wdr23KO* mice liver tissue (**Figure 2A**). IDE is a major enzyme responsible for insulin degradation that plays a central role in hepatic glucose metabolism [26]. As such, the increased levels of IDE could contribute to the change in insulin sensitivity observed. To support this finding, we subsequently confirmed the increased steady-state levels of IDE in fresh liver samples from age-matched WT and *Wdr23KO* mice by western blot analysis (**Figure 2B-D**).

**Figure 2.**
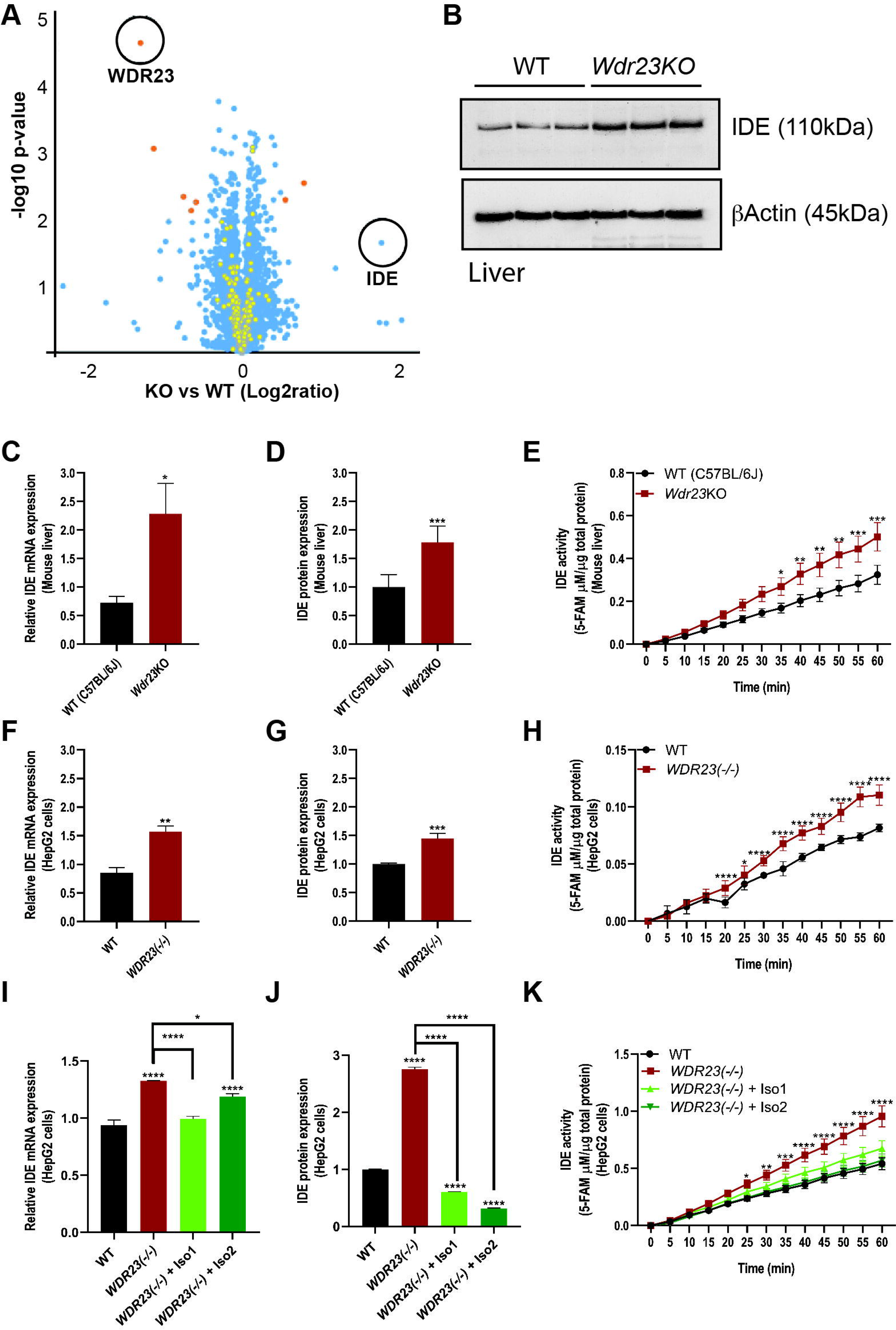
Loss of *Wdr23* increases insulin-degrading enzyme (IDE) activity in the liver. An unbiased proteomic assessment of proteins in liver from WT and *Wdr23KO* animals reveals an increase in IDE **(A)**, which is confirmed biochemically in freshly isolated livers (**B**). Increased IDE expression is increased at the mRNA (**C**) and protein (**D**) levels that results in enhanced enzymatic activity (**E**) in *Wdr23KO* livers, which is recapitulated in a HepG2 cell line with all copies of *WDR23* deleted “*Wdr23(-/-)*”(**F-H**). Rescue of *WDR23* isoforms suppresses the increased expression of IDE mRNA (**I**) and protein (**J**) and enhanced enzymatic activity (**K**) although IDE mRNA remain high in cells rescued for *WDR23* isoform 2 as compared to WT. *p<.05, **p<.01, ***p<.001, ****p<.0001

In addition, we assessed IDE proteolytic activity, which was significantly enhanced in liver homogenates from *Wdr23KO* mice as compared to WT (**Figure 2E**). A similar enhancement of IDE enzymatic activity was measured in hepatocytes isolated from *Wdr23KO* mice as compared to WT hepatocytes (Figure S2A-D) suggesting a cellular defect in the major parenchymal cells of the liver [27]. To further confirm specificity of the WDR23-dependent regulation of IDE, we developed a HepG2 cell line where we deleted all copies of *WDR23* by CRISPR/Cas9 genomic editing, hereafter called *WDR23(-/-)* (Figure S2E). As we observed in isolated liver tissues and primary hepatocytes, *WDR23(-/-)* HepG2 cells exhibit an increased level of steady-state IDE expression and enzymatic activity (**Figure 2F-H** and Figure S2F). Importantly, transfection of HepG2 cells with GFP:*WDR23* expression plasmids [17, 19] abolished the increased IDE expression and enhanced enzymatic activity (**Figure 2I-K** and Figure S2G). These data reveal that IDE expression and activity is linked to the insulin metabolism defects in *Wdr23KO* mice.

### Loss of *Wdr23* interferes with the expression of glucose metabolism pathway

Based on the altered transcriptional levels of several metabolic homeostasis genes, we next performed next generation RNA-sequencing analyses to discern the scope of transcripts that are sensitive to the activity of the WDR23 proteostasis pathway (**Figure 3A**). Significantly, pathway analysis using the GO and KEGG databases revealed the dysregulated genes induced by *Wdr23* deletion were enriched in carbohydrate metabolic processes that are regulated by insulin signaling and PPAR signaling pathways (**Figure 3B** and Table S1-2). These genes are influenced by insulin secretion and signaling cascades [28, 29], which were consistent with protein expression results from both isolated liver tissue and HepG2 cells. We further assessed the differentially expressed genes which revealed enrichment for components of the AGE/RAGE signaling pathway which regulates glucose metabolism in patients with diabetic complications [30] (**Figure 3B** and Table S2). Taken together, the loss of *Wdr23* alters transcription of glucose and insulin metabolism pathways that result in a shift in metabolic homeostasis.

**Figure 3.**
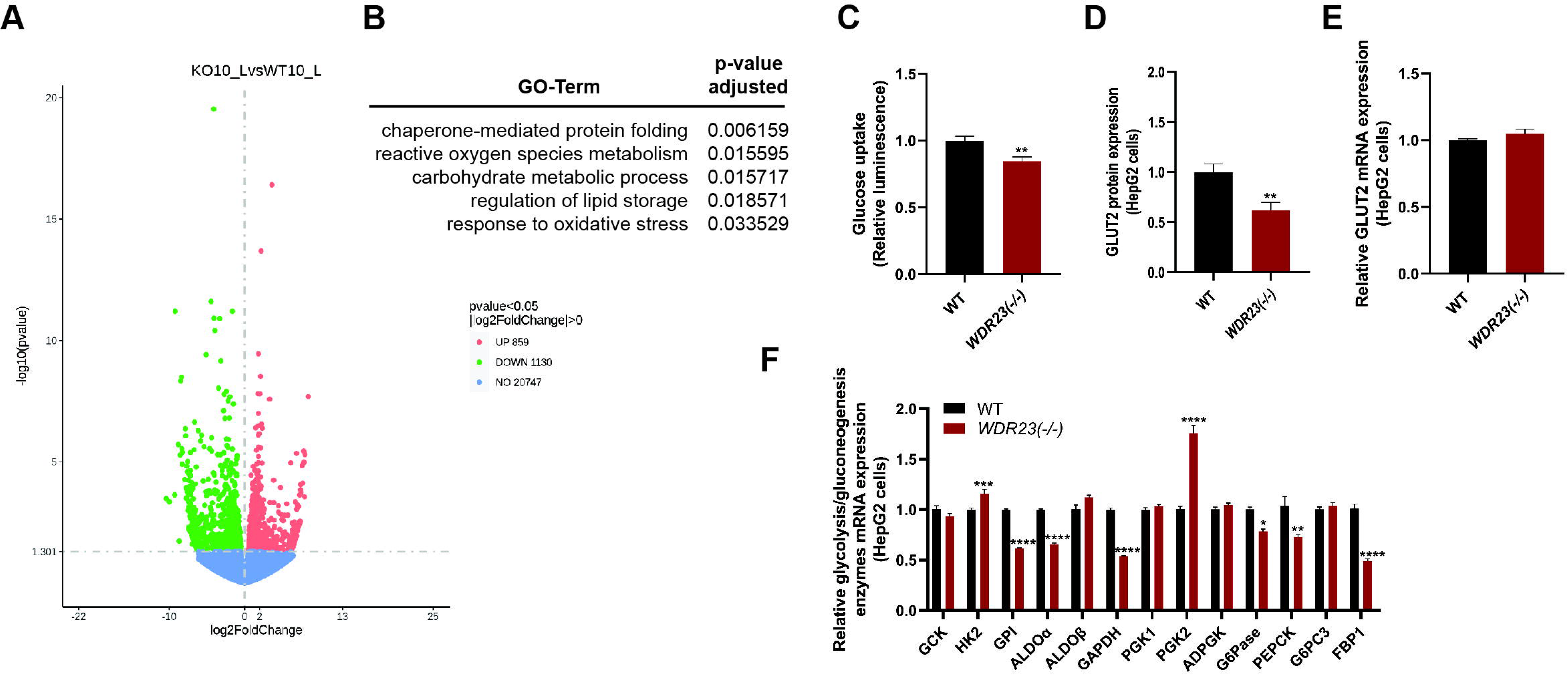
Loss of *Wdr23* disrupts glucose metabolism. RNA-sequencing analysis of male liver tissues isolated from *Wdr23KO* mice compared to WT (C57BL/6J). Volcano plot **(A)** and lists of all differentially expressed genes **(B)** in *Wdr23*KO livers. Up-regulated and down-regulated genes are indicated as red and green, respectively. *Wdr23(-/-)* HepG2 cells have decreased capacity for glucose uptake (**C**) that correlated with reduced abundance of the major glucose transporter of GLUT2 protein (**D**) and mRNA (**E**), and dysregulated expression of glycolytic and gluconeogenic enzymes (**F**). *p<.05, **p<.01, ***p<.001, ****p<.0001

Glucose uptake by hepatocytes plays an important role in the liver metabolic homeostat and its response to insulin. To assess whether WDR23 contributes to intracellular glucose influx, we next examined glucose absorption in HepG2 cells with and without *WDR23*. Glucose uptake was significantly decreased in *WDR23(-/-)* HepG2 cells as compared to control HepG2 cells (**Figure 3C**). The major glucose transporter in the plasma membrane of hepatocytes is GLUT2 [31]. We measured the abundance of GLUT2 mRNA (**Figure 3E**) and protein (**Figure 3D)** and (Figure S3A) which was significantly reduced in *WDR23(-/-)* HepG2 cells and is consistent with the reduced capacity of these cells to transport extracellular glucose. Intriguingly, SLC2A8/GLUT8, which can transport trehalose, a disaccharide consisting of two glucose molecules [32], was identified as ∼2-fold upregulated in liver tissues from the *Wdr23KO* mice that may represent a compensatory response to increase carbohydrate influx (Table S4). In addition, the galactose metabolism enzyme GALE was enriched in *Wdr23KO* livers which could aid in the utilization of carbohydrate alternatives to glucose to meet cellular metabolic demands.

Based on the impaired ability for *WDR23(-/-)* HepG2 cells to transport glucose, we next measured the expression of several key enzymes in the cellular glycolysis and gluconeogenesis pathway. Consistent with a defect in glucose availability, the mRNA expression levels of glucose pathway enzymes including gluconeogenesis pathway genes: Glucose 6-phosphatase (G6Pase), Phosphoenolpyruvate carboxykinase (PEPCK), and Fructose-1,6-bisphosphatase (FBP1); and glycolysis pathway genes: Hexokinase 2 (HK2), Glucose-6-phosphate isomerase (GPI), Fructose-bisphosphate aldolase alpha (ALDOα), Glyceraldehyde 3-phosphate dehydrogenase (GAPDH), and Phosphoglycerate kinase 2 (PGK2), were significantly changed in *WDR23(-/-)* samples when compared with WT controls (**Figure 3F**). Taken together, these data reveal a significant change in the metabolic state of hepatic cells in response to the loss of *Wdr23*.

### Loss of *Wdr23* mimics insulin treatment

The insulin signaling cascade begins with insulin hormones that bind to the insulin receptor (IR), which then trigger the activation of two major kinase-dependent phosphorylation cascades through IRS1/PI3K/AKT and Ras/MAPK pathways [33]. Although there was a significant decrease in the level of circulating insulin in the *Wdr23KO* mice, which is consistent with an increased level of IDE expression that degrades insulin, we noted an increase in the phosphorylation state of several key mediators of the insulin signaling pathway (**Figure 4A**). We could not detect a significant change in the phosphorylation state of IRS-1 (**Figure 4B** and Figure S3B), which is phosphorylated in response to insulin binding at the insulin receptor, but phosphorylation of AKT2 in *WDR23(-/-)* HepG2 cells was increased >3-fold (**Figure 4C** and Figure S3C) and approximately 1.5-fold in liver homogenates from *Wdr23KO* mice (Figure S4A). Similarly, phosphorylation of MAPK was increased >2-fold in *WDR23(-/-)* HepG2 cells (**Figure 4D** and Figure S3D) and ∼1.5-fold increased *Wdr23KO* liver (Figure S4B). The PI3K/AKT axis of the insulin signaling cascade regulates metabolic homeostasis through multiple downstream pathways, including the forkhead transcription factor family (FoxO) and the target of rapamycin (mTOR) [29]. In *WDR23(-/-)* HepG2 cells We found a 50% decrease in the phosphorylation state of mTOR (**Figure 4E** and Figure S3E) and a modest increase in the phosphorylation state of FoxO1 in *WDR23(-/-)* HepG2 cells (**Figure 4F** and Figure S3F), which was similar in liver homogenates from *Wdr23KO* mice for mTOR, but not FoxO1 (Figure S4C-D) . We next tested whether *WDR23(-/-)* HepG2 cells are sensitive to exogenous insulin treatment. With the exception of MAPK and mTOR phosphorylation, *WDR23(-/-)* HepG2 cells mimic the phosphorylation state of WT cells treated with insulin and treatment of *WDR23(-/-)* HepG2 cells with insulin further enhances insulin cascade phosphorylation (**Figure 4A-F** and Figure S3B-F).

**Figure 4.**
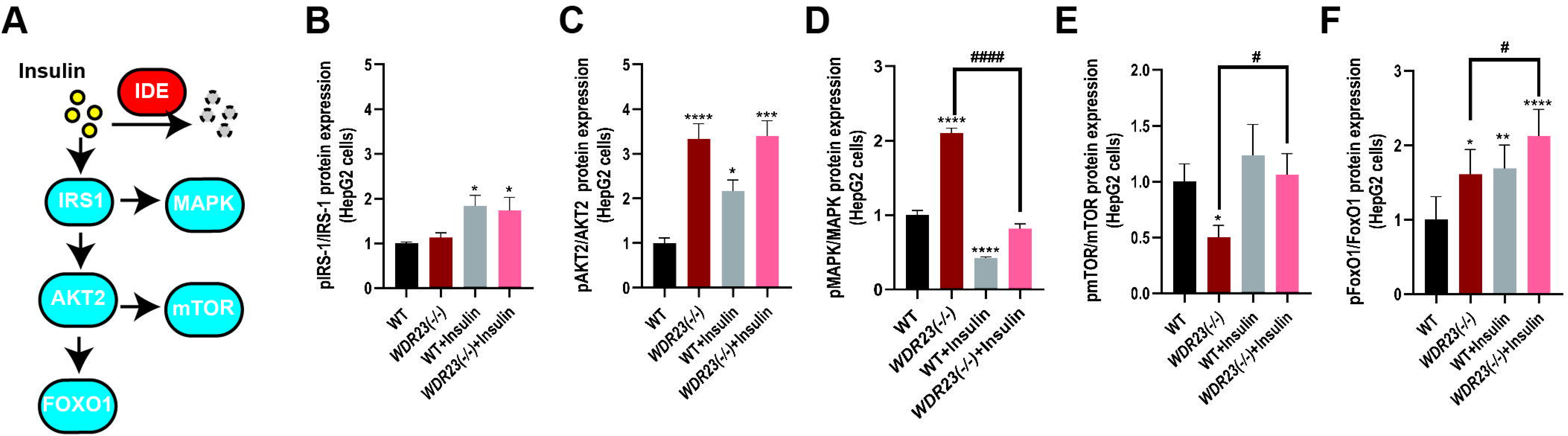
Loss of WDR23 disrupts insulin signaling. The effects of loss of *WDR2*3 in HepG2 cells, *WDR23(-/-),* on the insulin signaling pathway **(A)** as measured by the phosphorylation state of (**B**) IRS1 (Tyr608/Tyr612), (**C**) AKT2 (Ser474), (**D**) MAPK (Thr202/Tyr204), **(E)** mTOR (Ser2448), and (**F**) FoxO1 (Ser256). HepG2 cells were treated with 500 nM insulin and 25 mM glucose for 24h (+Insulin). *p<.05, **p<.01, ***p<.001, ****p<.0001, compared to WT. ^#^p<.05, ^##^p<.01, ^###^p<.001, ^####^p<.0001, compared to *WDR23(-/-)* HepG2 cells.

To link the changes in IDE expression with the changes in the phosphorylation of the insulin signaling pathway, we made use of ML345, an documented chemical inhibitor of IDE activity [34]. We first established the dose of ML345 that could effectively inhibit 50% activity in both WT and *WDR23(-/-)* HepG2 cells (Figure S5A) and subsequently treated cells with this concentration of inhibitor and measured the phosphorylation status of the insulin signaling pathway proteins (**Figure 5A-F**). Although inhibition of IDE activity had minimal effects on IRS1 phosphorylation (**Figure 5B** and Figure S5B), ML345 treated did reverse the enhanced phosphorylation of AKT2 (**Figure 5C** and **Figure S5C**), MAPK (**Figure 5D** and Figure S5D), and FoxO1 (**Figure 5F** and Figure S5F), further decreased the phosphorylation of mTOR (**Figure 5E** and Figure S5E). Taken together, these data support a model where the increased expression and activity of IDE in cells lacking WDR23 is causal for the defects in insulin responsiveness.

**Figure 5.**
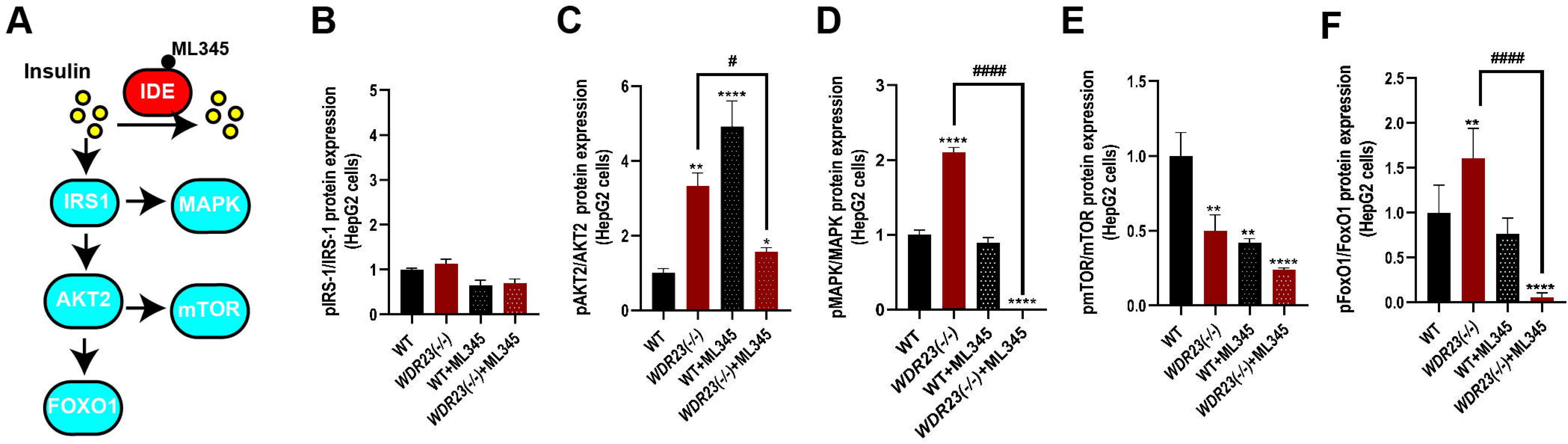
ML345 treatment reverses the effects of WDR23 loss on insulin signaling. The effects of treating WT and *WDR23(-/-)* HepG2 cells with the IDE inhibitor ML345 (40 µM, 24h) on the insulin signaling pathway as measured by the phosphorylation state of (**B**) IRS1 (Tyr608/Tyr612), (**C**) AKT2 (Ser474), (**D**) MAPK (Thr202/Tyr204), **(E)** mTOR (Ser2448), and **(F)** FoxO1 (Ser256). HepG2 cells were treated with 500 nM insulin and 25 mM glucose for 24h (+Insulin). *p<.05, **p<.01, ***p<.001, ****p<.0001, compared to WT. ^#^p<.05, ^##^p<.01, ^###^p<.001, ^####^p<.0001, compared to *WDR23(-/-)* HepG2 cells.

### Enhanced expression of IDE is mediated by the WDR23 substrate NRF2

We utilized ChEA3 for transcription factor enrichment analysis by orthogonal -omics integration [35] on our RNAseq data sets to identify transcription factors that mediate the responses to loss of *Wdr23* (Table S4). ChEA3 analysis revealed an enrichment for several transcription factors for the 309 significantly upregulated transcripts, including: CEBPB, which plays a significant role in adipogenesis, as well as in the gluconeogenic pathway; CREB3L3 that plays a crucial role in the regulation of triglyceride metabolism and is required for the maintenance of normal plasma triglyceride concentrations, and NFE2L2/NRF2, which was expected, as our previous work identified NRF2 as a direct target substrate of WDR23 [17, 19]. NRF2 is a conserved cytoprotective transcription factor that controls the expression of stress response and intermediary metabolism gene targets [36–38].

We were curious to test whether the changes in IDE expression and the subsequent insulin metabolism phenotypes in response to loss of *Wdr23* were associated with NRF2 activation. We searched *in silico* within the promoter region of *Ide* for the core ARE-like NRF2 consensus binding sequence [39], 5′-TGAC-3′, and found three putative binding sites for NFE2L2/NRF2 upstream of the translational start site for the human *IDE* locus and six ARE core elements in the mouse genome {29087512;29126285;27924024} (**Figure 6A**).

**Figure 6.**
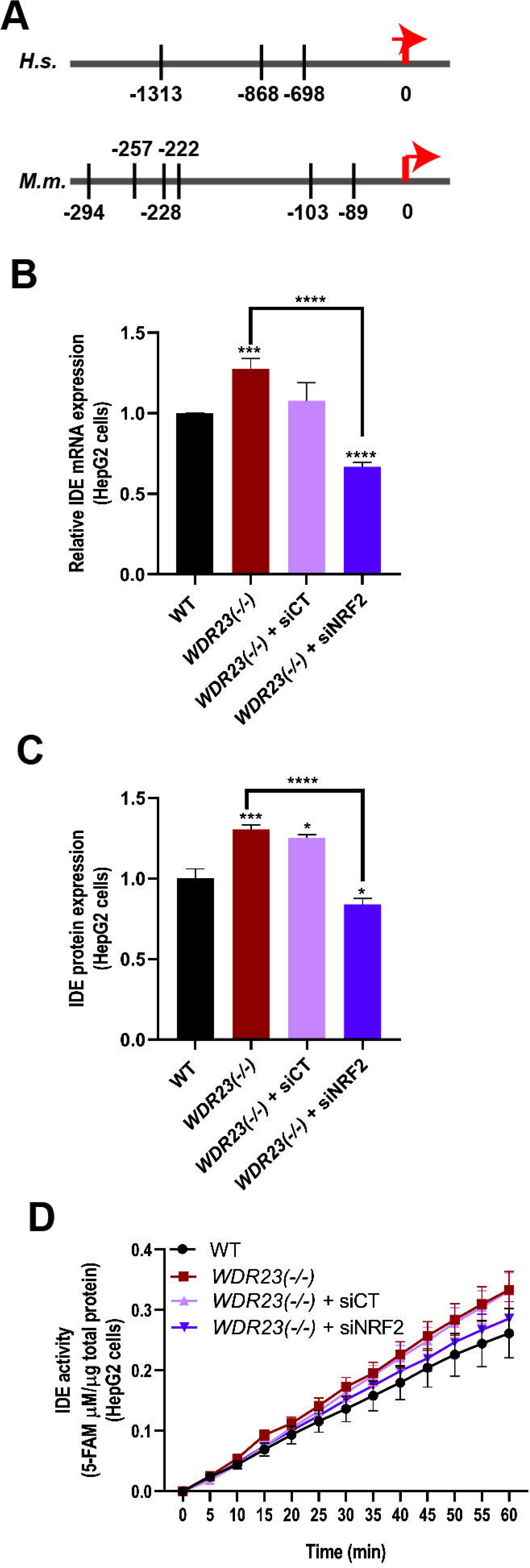
NRF2 mediates the Enhanced IDE activity in response to loss of *Wdr23*. Putative ARE core binding sites in promoter element of the human and mouse *IDE* gene (**A**); numbers are nucleotide position relative to translational start codon (0). The cytoprotective transcription factors NRF2 is necessary for the increased expression of IDE mRNA (**B**), protein (**C**), and insulin degradation enzymatic activity (**D**).

To confirm whether NRF2 is required for the increased expression of IDE in cells lacking *WDR23*, we examined the expression level and activity of IDE in WT and *WDR23(-/-)* HepG2 cells treated with *Nrf2*-specific siRNAs (Figure S6A). Following transfection of *NRF2-*specific siRNA, but not transfection with a scrambled siRNA, the enhanced expression of *IDE* transcripts (**Figure 6B**), IDE protein (**Figure 6C** and Figure S6B), and the enzymatic activity of IDE (**Figure 6D**) were abrogated. These data suggest that NRF2 transcriptional activity largely contributes to the changes in insulin-degrading enzyme when the WDR23 proteostasis pathway is impaired.

### Genetic variation in *Wdr23* is associated with the incidence of diabetes in older adults

Based on the remarkable conservation in the players and responses to altered insulin signaling in metabolic health, we were curious if human *WDR23* was associated with metabolic disease states. The US Health and Retirement Study (HRS) is a nationally representative survey of adults aged 50 years and older that is an innovative tool for investigating the normal aging processes [40–42]. Recently, the HRS data sets incorporated genotypic data of participants that enables the testing of associations between normal aging phenotypes and variation across genes [43, 44].

We assessed the available phenotypic data in the US Health and Retirement Study (HRS) for SNP associations with blood-based biomarkers of diabetes and found that genetic variation in *WDR23* is associated with altered hemoglobin A1C (HbA1c) levels. HbA1c is the standard biomarker measure used for the clinical diagnosis of diabetes and pre-diabetes [45]. In HRS, multivariable linear regression models were run to test for the association between each of the five annotated *WDR23* SNPs in the HRS datasets and hemoglobin A1c measurements for each individual **(Table 1)**, adjusting for age, gender, and principal components using PLINK. To correct for correlations between SNPs within the gene, we performed 1,000 permutations, comparing shuffled (null) data to the non-shuffled data to derive the empirical p-value threshold of 0.0142 for HbA1c for determining statistically significant associations with *WDR23* SNPs. Taken together, our study reveals that *WDR23* genotype (SNPs and genetic ablation) influences insulin metabolism and has the potential to impact human health during natural aging.

**Table 1.** Genetic variation in WDR23 is associated with age-related incidence of diabetes. HRS data association of *WDR23* variants reveals one variant with significant association with age-related diabetes as measured by HbA1c. Scan adjusted for age in 2006, sex, and 4 ancestral principal components. *p<.05; #p<.10 but considered suggestive with multiple test correction set at p=.01.

| SNP | Type | Position | Ref allele | NMISS | BETA | P | Significance |
| --- | --- | --- | --- | --- | --- | --- | --- |
| rs3742499 | Intron variant | 24586833 | A | 9326 | -0.015 | 0.1350 |  |
| rs2277481 | Intron variant | 24587545 | G | 9331 | -0.026 | 0.0129 | * |
| rs17101367 (kgp6424105) | Coding variant | 24587667 | A | 9328 | -0.012 | 0.2406 |  |
| rs2277482 | Intron variant | 24587795 | A | 9329 | -0.019 | 0.0670 | # |
| rs2277483 | Intron variant | 24591892 | G | 9333 | -0.019 | 0.0650 | # |

\* $p < .05$ ; # $p < .10$ all considered suggestive with multiple test correction set at $p = .01$

## DISCUSSION

In the present study, we reveal a role for WDR23 in the expression of IDE which suggests that targeting WDR23 could be a new therapeutic strategy for regulating insulin sensitivity. Several lines of evidence supported the role of WDR23 in insulin sensitivity. First, loss of WDR23 disrupts insulin sensitivity in mice. Second, livers from *Wdr23*KO mice have increased IDE expression and activity. Third, the transcriptional profiling and pathway analysis indicated that genes involved in glucose and lipid metabolism, especially insulin signaling pathways were dysregulated when *Wdr23* is absent.

IDE is a metalloendopeptidase with a high affinity for insulin which is a major enzyme responsible for hepatic insulin clearance [26, 46]. In addition to insulin, IDE degrades glucagon, beta-amyloid peptide, and atrial natriuretic peptide [47, 48]. Impaired insulin clearance is associated with lower IDE levels that are observed in T2DM patients [26, 46]. In addition, SNPs in *Ide* locus have been linked to the risk of T2DM [49], which is similar to our finding that SNPs in *Wdr23* are associated with the risk of diabetes at older age (**Table 1**). Here we demonstrate that liver tissues and hepatocytes from *Wdr23KO* mice and engineered HepG2 cells lacking WDR23 have induced IDE expression and activity with a parallel decrease in circulating plasma insulin levels (**Figure 1**). Correspondingly, *Wdr23KO* male mice display impaired insulin sensitivity, suggesting WDR23 is the essential regulator of insulin signaling mediated in part by the regulation of the IDE. However, there are no significant differences measured in glucose clearance and insulin tolerance in female *Wdr23*KO mice as compared to WT (**Figure S1**). These results consistent with human study by Hong et al., that the variants of gene encoding IDE are strongly associated with insulin clearance in men [50]. The insulin-dependent regulation of hepatic glucose and lipid metabolism is essential for metabolic homeostasis [51] and as such future investigation of the impact of *Wdr23* loss on lipid metabolism and hepatic steatosis, particularly on high fat diets will be of great interest.

An imbalance in insulin signaling can drive metabolic disease due to its activity as a regulator of cellular metabolic homeostasis [29]. The insulin signaling cascade begins with insulin binding to the insulin receptor (IR), which then trigger the activation of two major kinase-dependent phosphorylation cascades through IRS1/PI3K/AKT and Ras/MAPK pathways [33]. Our unbiased proteomic analyses further revealed significant changes in key downstream mediators of the transcriptional response to insulin signaling that are central to the maintenance of metabolic homeostasis.

AKT2 phosphorylation was increased in *WDR23(-/-)* HepG2 cells similar to WT cells treated with insulin (**Figure 4)**, but insulin treatment failed to further increase AKT2 phosphorylation suggesting an impairment of the insulin response in cells lacking *Wdr23*. Importantly, treatment with the IDE inhibitor ML345 reversed the AKT2 phosphorylation state in *WDR23(-/-)* HepG2 cells (**Figure 5**), indicating the causality of increased IDE in this response. Similarly, PI3K/AKT signaling regulates metabolic homeostasis through multiple downstream pathways, including the FoxO and the mTOR [33]. We noted a modest increase in the steady-state phosphorylation of FoxO1 in *WDR23(-/-)* HepG2 cells (**Figure 4**), while decrease in *Wdr23*KO liver (**Figure S4**). Although the culturing conditions of the *in vitro* cell model likely play a role here, beyond insulin, IDE has a low affinity for glucagon and amylin which may account for a part of this observed difference [47, 48]. Glucagon regulates hepatic gluconeogenesis by promoting phosphorylation of FoxO1 and increasing expression of genes encoding the rate-limiting enzymes responsible for gluconeogenesis such as G6Pase, PEPCK, FBP1, and glucose 6-phosphatase [52]. As such, the increase in IDE activity in *Wdr23*KO mice might affect both insulin and glucagon levels which synergistically influence FoxO1 phosphorylation to regulate gluconeogenesis. Beyond FoxO1, phosphorylated AKT can also inhibit mTORC1 signaling and the subsequent activation of SREBP-1 and fatty acid synthase and cholesterol-related genes [33]. The significant reduction of mTOR phosphorylation in both *WDR23(-/-)* HepG2 cells (**Figure 4**) and *Wdr23*KO liver (**Figure S4**) further supports the activation of AKT2 signaling and reveals the importance of future work to investigate the impact of loss of *WDR23* on lipid metabolism.

The Keap1-NRF2 system is a critical target for preventing T2DM [37] as demonstrated by the activation of NRF2 through KEAP1 knockdown, which promotes glucose uptake and insulin sensitivity in diabetic mice [37, 53]. Moreover, NRF2 deletion impairs glucose handling, lipid peroxidation and cytoprotective gene induction in mice [54] as several enzymes and proteins involved in hepatic lipogenesis and gluconeogenesis are encoded by ARE-containing NRF2 target genes [55]. Our previous work has demonstrated that WDR23 is second mechanism for regulating NRF2 activity [17, 19], similar to the role of *C. elegans* WDR-23 on SKN-1 [17, 56–58]. Our results confirm that the increase in IDE expression in the absence of WDR23 is dependent on NRF2. As such, our study reveals a new regulatory axis of insulin homeostasis mediated by WDR23-CUL4 regulation of NRF2 and subsequent activation of IDE. Although the connection between the WDR23-mediated regulation of NRF2 by the ubiquitin-proteasome system is promising, the connection to IDE and diabetes requires further study to clarify how other WDR23 targets might influence metabolic homeostasis.

Although several WDR23 SNPs were found to have a significant association of HbA1c in older adults of the HRS we cannot exclude the possibility that these SNPs mark additional tightly linked genes in this region. However, the similarities observed in our mouse model and HepG2 cell lines suggests that WDR23 genotype is important across mammals. In the future, as the HRS genotypic data expands and becomes more diverse, an assessment of whether sex and ethnicity are significant drivers of the association of *WDR23* genotype and diabetes will be of great interest.

Collectively, our study suggests that *WDR23* plays an important role in insulin signaling and metabolic homeostasis and can provide an important data point in the development of a personalized medicine approach to ensure optimal health with age.

## ACKNOWLEDGEMENTS

We thank J Gonzalez, M Donoghue, M Lynn for mouse colony assistance, and Dr. R Irwin, Dr. N Stuhr, B Van Camp and M Ramos for critical reading of the manuscript. This work was funded by the NIH RF1 AG063947 and R01 AG058610 to SPC and a Glenn Foundation for Medical Research Postdoctoral Fellowship in Aging Research from the American Federation for Aging Research to CD. This study was supported in part by funding from The National Institute on Aging, through the USC-Buck Nathan Shock Center (P30 AG068345). The National Institute on Aging has supported the collection of both survey and genotype data for the Health and Retirement Study through co-operative agreement U01 AG009740. The datasets are produced by the University of Michigan, Ann Arbor. The HRS phenotypic data files are public use datasets, available through: https://hrs.isr.umich.edu/data-products/access-to-public-data. The HRS genotype data are available to authorized researchers: https://www.ncbi.nlm.nih.gov/projects/gap/cgi-bin/study.cgi?study_id=phs000428.v2.p2https://www.ncbi.nlm.nih.gov/projects/gap/cgi-bin/study.cgi?study_id=phs000428.v2.p2. We thank the Wellcome Trust Sanger Institute Mouse Genetics Project (Sanger MGP) and its funders for providing the *Wdr23(tm1a)* mutant mouse line.

## AUTHOR CONTRIBUTIONS

Conceptualization, S.P.C.; methodology, T.E.A. and S.P.C.; formal analysis, C.D., B.N.S., T.E.A. and S.P.C.; investigation, C.D., B.N.S., and T.E.A., and S.P.C.; data curation, C.D., B.N.S., T.E.A. and S.P.C.; writing – original draft, C.D. and S.P.C.; writing – review & editing, S.P.C.; visualization, C.D., T.E.A., and S.P.C.; supervision, S.P.C.; project administration, S.P.C.; funding acquisition, C.D. and S.P.C.

## SUPPLEMENTAL FIGURE LEGENDS

**Figure S1. Assessment of female *Wdr23*KO mice.**

Female *Wdr23*KO mice gain weight at the same rate as age-matched C57BL/6J controls (**A,B**). Glucose tolerance, as measured by glucose tolerance test (GTT) is similar to C57BL/6J control animals at 10-weeks (**C,D**) and 1-year of age (**E,F**). Insulin tolerance, as measured by insulin tolerance test (ITT) is similar in Wdr23KO as compared to C57BL/6J animals when fed a standard chow (**G,H**) or 10% fat (**I,J**) diet. Analysis of fat deposits in the liver of C57BL/6J animals (**K-M**) is unremarkable as compared to age-matched *Wdr23*KO animals (**N-P**).

**Figure S2. IDE expression and activity is enhanced in hepatocytes from *Wdr23KO* liver.**

Hepatocytes derived from isolated liver tissue of *Wdr23KO* animals have higher expression of IDE protein (**A,C**), mRNA (**B**) that results in an increase in relative enzymatic activity (**D**). Generation of a HepG2 cell line with deletion of *Wdr23*, “*WDR23(-/-)*”, results in undetectable full length WDR23 from gDNA and significant >90% reduction in full length mRNA (**E**). *WDR23(-/-)* HepG2 cells display increased IDE protein (**F**) that is reduced with Wdr23-I or Wdr23-II are rescued (**G**).

**Figure S3. Loss of WDR23 disrupts homeostatic activation of the insulin signaling cascade.**

Loss of WDR23 results in decreased glucose transporter 2 protein expression (GLUT2) (**A**) Western blot analysis of the impact of loss of *WDR23* on the phosphorylation state of IRS1 (Tyr608/Tyr612) (**B**), AKT2 (Ser474) (**C**), MAPK (Thr202/Tyr204) (**D**), mTOR (Ser2448) (**E**) and FoxO1 (Ser256) (**F**) in *WDR23(-/-)* HepG2 cells. HepG2 cells (+Insulin) were treated with 500 nM insulin and 25 mM glucose for 24 h to induce insulin resistant.

**Figure S4. Loss of *Wdr23* disrupts the insulin signaling pathway in the liver of male mice.**

Phosphorylation of AKT2 (Ser474) (**A**) and MAPK (Thr202/Tyr204) (**B**) are increased, while phosphorylation mTOR (Ser2448) (**C**) and FoxO1 (Ser256) (**D**) are decreased in *Wdr23*KO mice liver sample when compared to age matched WT (C57BL/6J) control. Male mice were fed 10% fat diet for 44 weeks before liver tissue harvesting.

**Figure S5. Impact of IDE inhibition on insulin signaling cascade.**

ML345 inhibited IDE activity in WT and *WDR23 (-/-)* HepG2 cells (**A**). Western blot analysis of the impact of loss of *WDR23* on the phosphorylation state of IRS1 (Tyr608/Tyr612) (**B**), AKT2 (Ser474) (**C**), MAPK (Thr202/Tyr204) (**D**), mTOR (Ser2448) (**E**) and FoxO1 (Ser256) (**F**) in *WDR23(-/-)* HepG2 cells after treatment with IDE inhibitor (ML345). HepG2 cells were treated with 40 µM ML345 for 24 h to inhibit IDE activity. HepG2 cells (+Insulin) were treated with 500 nM insulin and 25 mM glucose for 24 h to induce insulin resistant.

**Figure S6. NRF2 regulates the transcription of IDE resulting from loss of WDR23.** NRF2 levels are significantly reduced in cells transfected with NRF2 siRNAs but not a scrambled siRNA sequence (**A**). The enhanced expression of IDE protein was abrogated in *WDR23(-/-)* HepG2 cells treated with NRF2-specific siRNA (**B**).

## SUPPLEMENTAL TABLE LEGENDS

**Table S1.** The GO functional enrichment analysis of DEGs in *Wdr23*KO mice liver tissues compare to the WT (C57BL/6J) control with the threshold of p≤0.05.

**Table S2.** The KEGG pathway enrichment analysis of DEGs in *Wdr23*KO mice liver tissues compare to the WT (C57BL/6J) control with the threshold of p≤0.05.

**Table S3.** Differential expression of targeted proteins in *Wdr23*KO mice liver tissues compare to the WT (C57BL/6J) control with the threshold of p≤0.05.

**Table S4.** ChIP-X Enrichment Analysis 3 (ChEA3) of transcription factors in *Wdr23*KO mice liver tissues compare to the WT (C57BL/6J) control with the threshold of p≤0.05.

## MATERIALS AND METHODS

### Key resources table

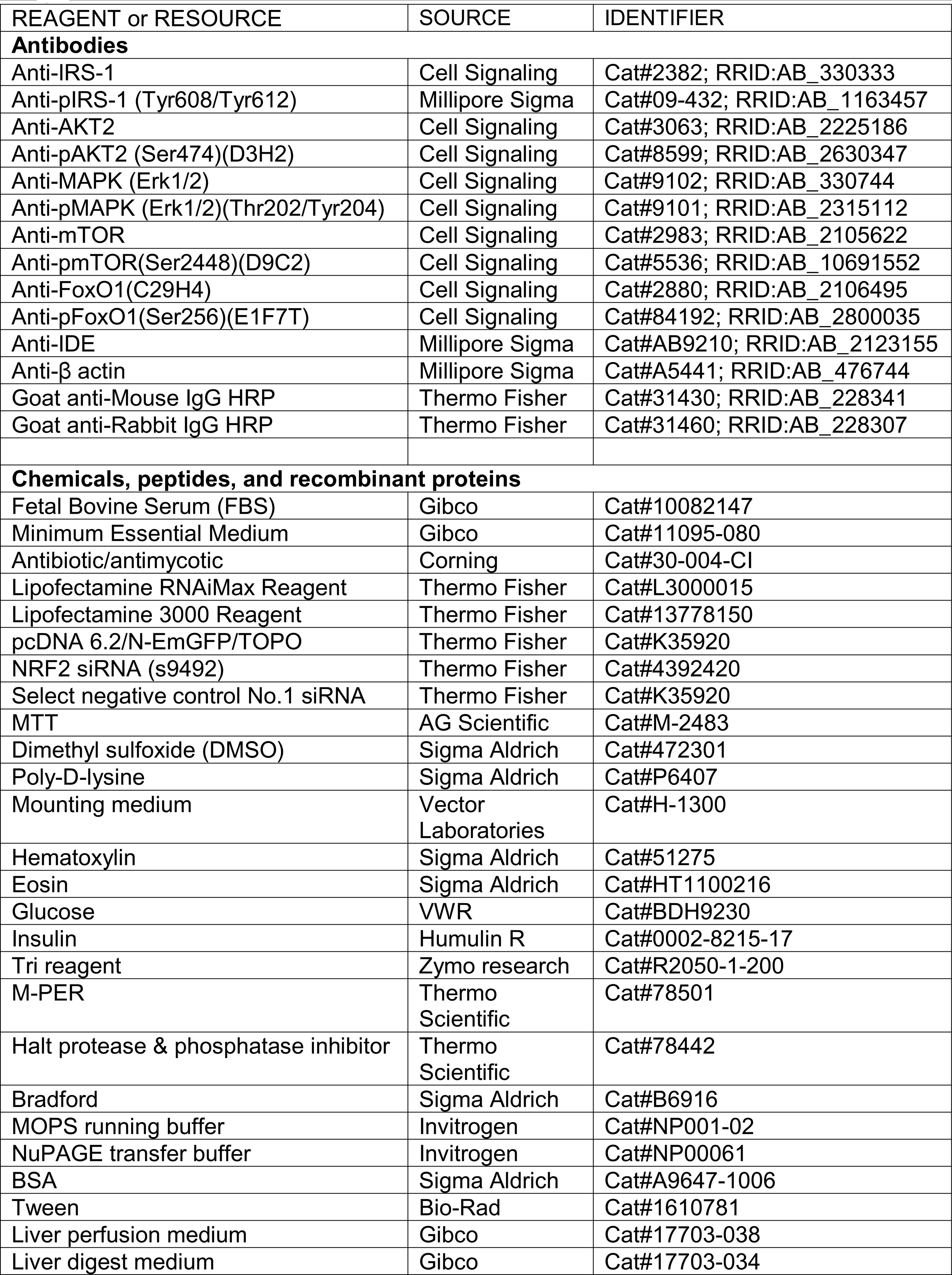
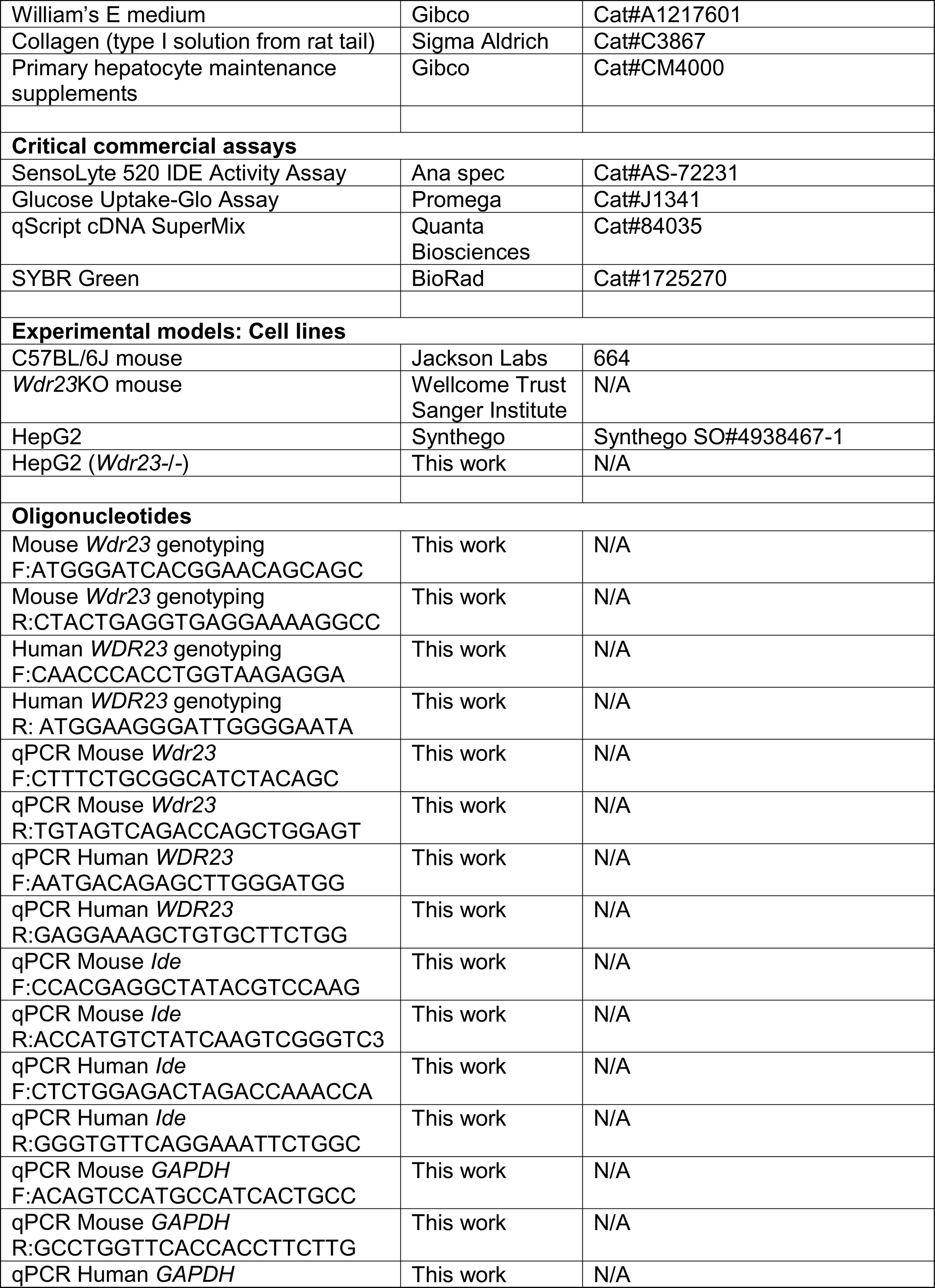
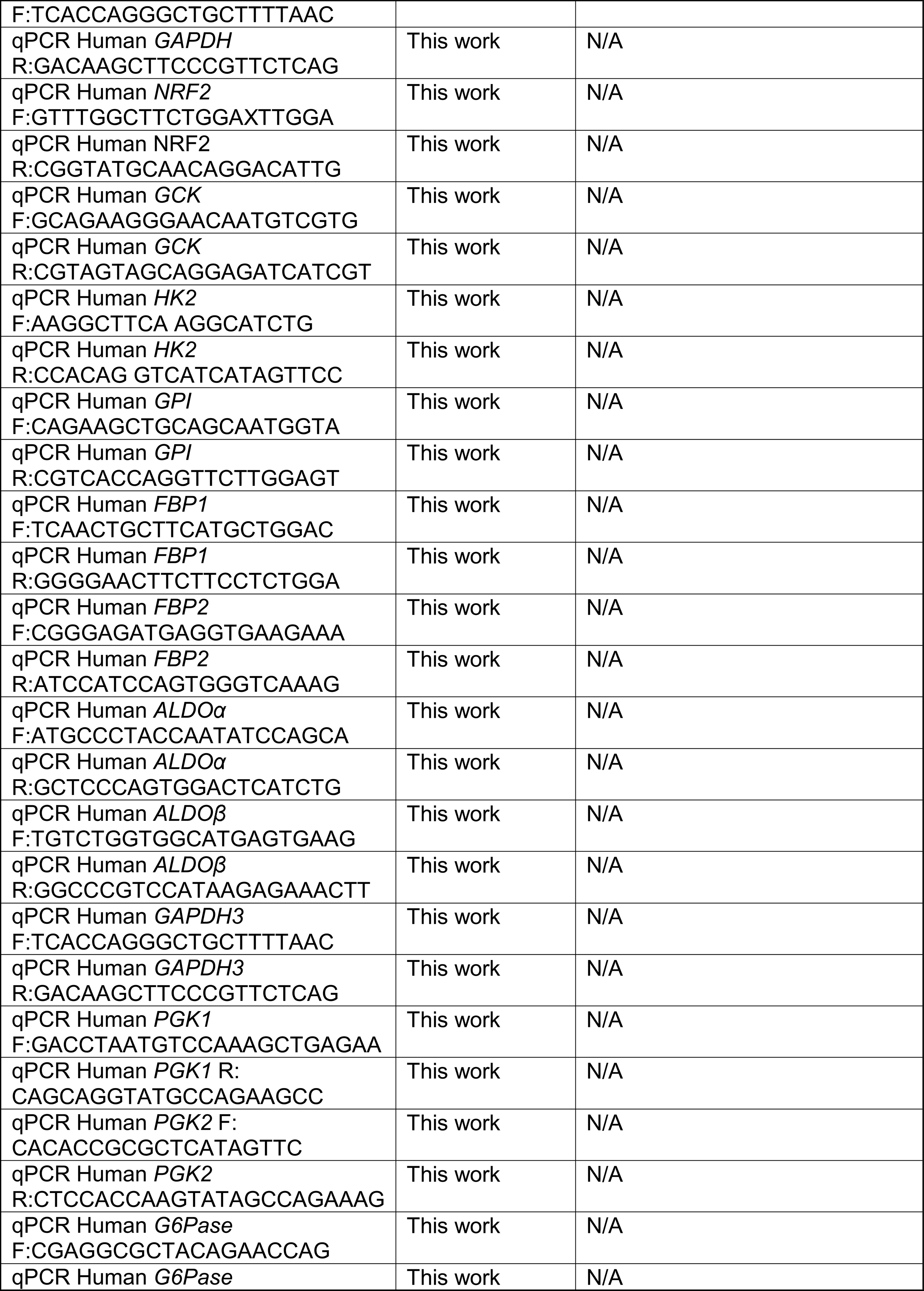
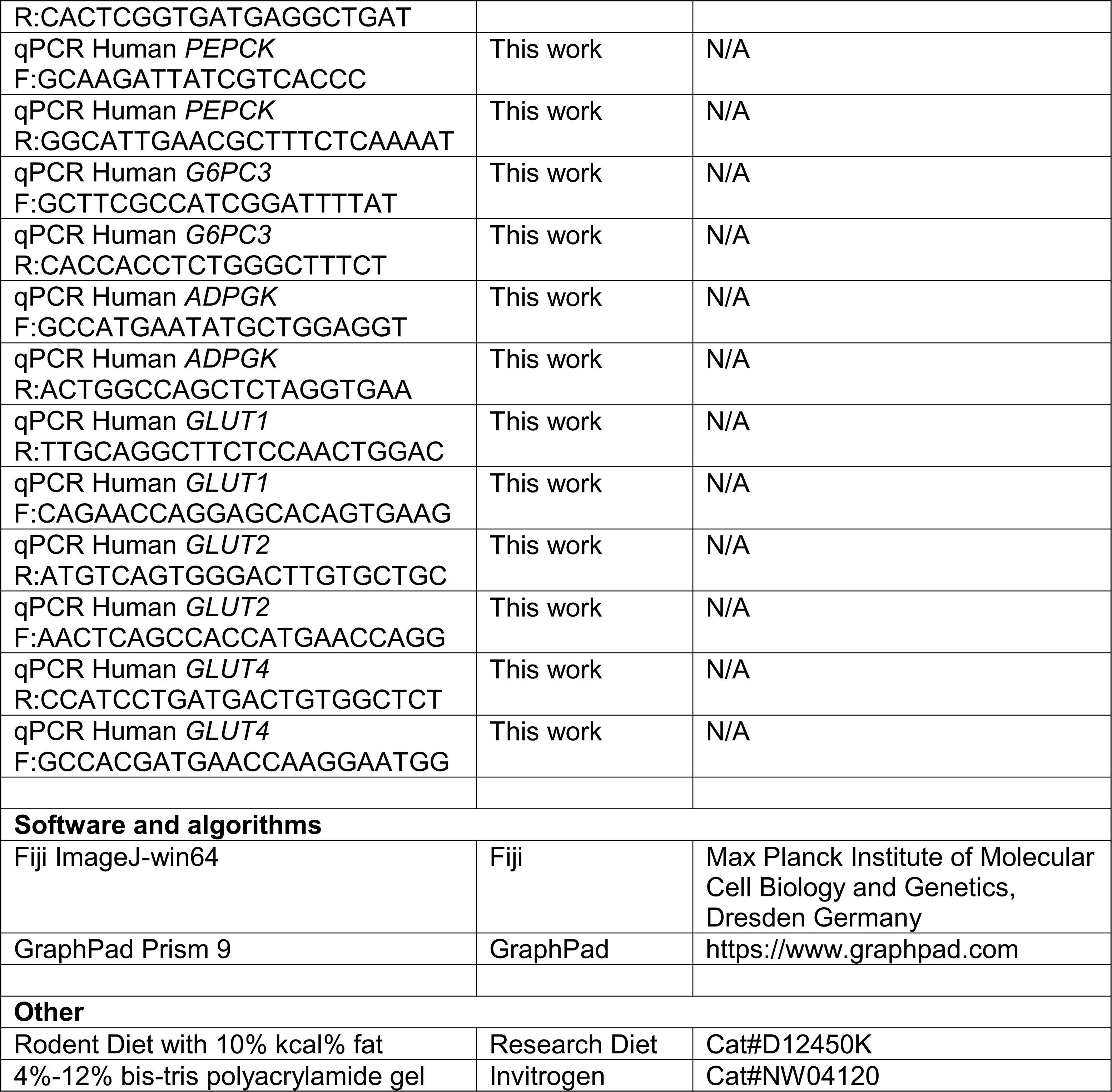

### Animals

All animal protocols were approved by the Institutional Animal Care and Use Committee (IACUC) of the University of Southern California and all the procedures were conducted in compliance with institutional guidelines and protocols.

*Wdr23* knock-out (*Wdr23*KO) mice were generated by Wellcome Trust Sanger Institute [59, 60]. *Wdr23KO* animals were subsequently backcrossed nine times into our C57BL/6J (WT) strain from the Jackson laboratory. Heterozygous (*Wdr23 +/-*) *dams* and *sires* were then mated to generate *Wdr23+/*+ and *Wdr23-/-* animals that were maintained as WT and KO, respectively. Male and female mice (n=4-6/group) were kept in a 12:12 h light-dark cycle, constant temperature, and humidity room. All animals were allowed *ad libitum* access to water and food.

### Rodent diets

Mice were fed ad lib with irradiated D12450K rodent chow (Research Diets), containing 16.1kJ/g of digestible energy (animal-based protein 3.22kJ/g, carbohydrate 11.27kJ/g, fat 1.61 kJ/g).

### Glucose tolerance tests (GTT)

Performed as previously described [61], animals (10 weeks and 1 year old) were fasted for 18 h prior to administration of glucose (2g/kg bodyweight) via intraperitoneal injection. Blood glucose was measured from the tail tip at 0, 15, 30, 60-, 90-, 120- and 180-min post-injection (Contour Next One Blood Glucose monitoring (9763).

### Insulin tolerance tests (ITT)

Performed as previously described [61], animals (30- and 40-weeks old mice) were fasted for 4 hours and then injected intraperitoneally with recombinant human insulin at (0.5 U/kg bodyweight) and glucose levels were determined at 0, 15, 30, 60, 90, 120 and 180 min after insulin injection.

### IDE activity assay

The enzymatic activity of IDE in mice liver tissue and HepG2 cells were determined using Sensolyte 520 IDE Activity Assay fluorometric kit according to manufacturer’s protocol.

### Determination of plasma hormones

Animals were anesthetized, blood samples were taken and centrifuged for 10 min at 10000 rpm, and the supernatant (plasma) was used for hormone measurement. Plasma insulin levels were measured using an ELISA kit (Meso Scale Discovery) according to manufacturer’s instructions.

### RNA extraction and real-time quantitative PCR

Quantitative PCR was performed as previously described [19]. Briefly, mice liver tissue or HepG2 cells were collected and lysed in Tri reagent (Zymo research). RNA was extracted according to the manufacturer’s protocol. RNA was reverse-transcribed to complementary DNA using qScript cDNA SuperMix (Quanta Biosciences). Quantitative PCR was conducted by using SYBR Green (BioRad). The relative expression of each gene was normalized against the internal control gene (GAPDH), and expression levels were analyzed using the 2-ΔΔCT method. The gene-specific sequences of the primers for HepG2 cells and mouse liver tissue were represented in key resources table.

### RNA-seq

Isolated RNA was sent to Novogene for library preparation and deep sequencing in biological triplicate. The read counts were used for differential expression (DE) analysis by using the R package DEseq2 (R version 3.5.2). Differentiated expressed genes were analyzed using p value <0.05 and fold change >1.5 as cutoff.

### Western blot analysis

Whole cell lysates were prepared in M-PER buffer (1x Mammalian Protein Extraction Reagent (Thermo Scientific), 0.1% Halt Protease & Phosphatase inhibitor (Thermo Scientific) according to the manufacturer’s protocol. Total protein concentrations were quantified by Bradford assay (Sigma). An equal amount of protein (20 µg) was separate on 4%-12% bis-tris polyacrylamide gel (Invitrogen) in MOPS running buffer (Invitrogen) and then transferred to nitrocellulose membranes (GE Healthcare Life science). After blocking for 1 h with 3% BSA in PBST (PBS, 0.1% Tween 20), the membranes were subjected to immunoblot analysis. Antibodies used include: IDE (Millipore sigma, 1:5000), IRS-1 (Cell Signaling Technology, 1:500), pIRS-1 (Millipore Sigma, 1:500), AKT (Cell Signaling Technology, 1:1000), pAKT (Cell Signaling Technology, 1:1000), AKT2 (Cell Signaling Technology, 1:1000), pAKT2 (Cell Signaling Technology, 1:1000), MAPK (Cell Signaling Technology, 1:1000), pMAPK (Cell Signaling Technology, 1:1000), mTOR (Cell Signaling Technology, 1:1000), pmTOR (Cell Signaling Technology, 1:1000), FoxO1 (Cell Signaling Technology, 1:1000), pFoxO1 (Cell Signaling Technology, 1:1000), β-actin (Millipore Sigma, 1:10000) and HRP-conjugated secondary antibodies (Thermo Fisher, 1:10,000). Specific protein bands were visualized and evaluated using FluorChem HD2 (ProteinSimple). The full images of electrophoretic blots were represented in supplementary materials.

### Isolation of mouse primary hepatocytes

Mouse primary hepatocytes were isolated from male *Wdr23*KO and WT mice, aged 3-4 weeks, using a modified collagenase perfusion methods. Briefly, the liver was perfused via the portal vein with perfusion medium (GIBCO) for 6 mins, and liver digest medium (GIBCO) for 5 mins. Liver was removed and placed in a 100 mm plate filled with cold washing medium (Williams’ E medium (WEM, GIBCO), supplemented with primary hepatocyte maintenance supplements (GIBCO). Liver was dispersed into small pieces in the medium using forceps and filtered through a 100/70 µM cell strainer into a falcon tube. Cells are collected by centrifugation at 50 g for 3 min and washed 3 times with 20 ml washing medium. The cells were counted, and viability was evaluated by trypan blue exclusion. Hepatocytes were plated in 6 well plates (pre-coated with 0.01% collagen in acetic acid (Sigma) at a density of 2x10^5^ cells per well in maintenance medium (WEM, GIBCO), supplemented with primary hepatocyte maintenance supplements (GIBCO) and incubated for 2-3 hours, 37°C, 5% CO_2_.

### Cell culture and transfections

WDR23 depleted (*WDR23(-/-)*) HepG2 cells were generated by CRISPR/Cas9 (Synthego). Cells were maintained in Minimum Essential Medium (GIBCO) supplemented with 10% fetal bovine serum (GIBCO) and 1% antibiotic/antimycotic (Corning) at 37°C, 5% CO_2_.

Full-length cDNA sequence of Hs *WDR23* Isoforms 1 and 2 were cloned into pcDNA 6.2/N-EmGFP/TOPO (Thermo Fisher), as previously described [19]. siRNAs (Thermo Fisher) used include: NRF2 (s9492) and control No.1 (4390843).

Transfections were performed with Lipofectamine 3000 (Thermo Fisher) and Lipofectamine RNAiMAX (Thermo Fisher) according to the manufacturer’s protocol. For establishment of insulin-resistance in HepG2 cells, cells were seeded in 6-well plates with normal medium for overnight. After reaching 80% confluence, the medium was replaced with SF-MEM and incubated for 24h. Subsequently, the cells were treated with SF-MEM supplemented with 500 nM insulin and 25 mM glucose for 24 h.

### Glucose uptake measurements

Glucose uptake was performed by using Glucose Uptake-Glo from Promega. Briefly, HepG2 cells were seeded in 96-well plates (2×10^4^ cells/well) for 24 hours. *WDR23(-/-)* HepG2 cells were transiently transfected with indicated plasmids for 24 hours. Samples were prepared in triplicates for Glucose Uptake-Glo according to manufacturer’s protocol, and another setup was for cell viability assay to normalize the cell number.

### Cell viability

MTT was used to measure cell viability. 0.5 mg/ml MTT were added to the culture medium and incubated for 3 h at 37°C. The formazan crystals were solubilized by DMSO.

The absorbance (550) or luminescence were measured using a SPECTRA max M2 Plate Reader. Intracellular glucose uptake was expressed as relative luminescence which normalized by cell viability.

### Microscopy

### Fluorescence-based imaging

HepG2 cells were grown on coverslips (coated with poly-D-lysine, Sigma) and transiently transfected with *WDR23*-isoform plasmids for 24 hours. The coverslips were mounted (Vector) on cover slides and imaged with a Zeiss Axio Imager.M2m microscope, Axio Cam MRm camera and Zen Blue software.

### Histological analysis

Liver sections were stained with hematoxylin and eosin (H&E) to visualize adipocytes and inflammatory cells in the tissues. Sections and cells were analyzed by Thunder Imaging Leica DMi8 microscope. H&E-stained sections (six slides for each sample) were randomly selected and quantified for steatosis area using Fiji ImageJ-win64 (Max Planck Institute of Molecular Cell Biology and Genetics, Dresden Germany).

### Protein mass spectroscopy

Proteomic characterization of the proteome of mice liver tissues were performed by Poochon Scientific. Briefly, the total protein extractions of tissue samples were prepared following Poochon SOP#602 protocols. The protein concentration of the supernatants was determined by BCA protein assay kit. 90 μg of protein lysate from each sample was run on SDS-PAGE followed by in-gel trypsin digestion, TMT-10plex labeling and LC/MS/MS analysis. The LC/MS/MS analysis was carried out using a Thermo Scientific Q-Exactive hybrid Quadrupole-Orbitrap Mass Spectrometer and Thermo Dionex UltiMate 3000 RSLCnano System. Each peptide fraction was load onto a peptide trap cartridge at a flow rate of 5 μl/min. The trapped peptides were eluted onto a reversed-phase 20 cm C18 PicoFrit column (New Objective, Woburn, MA) using a linear gradient of acetonitrile (3-36%) in 0.1% formic acid, for 100 min at a flow rate of 0.3 μl/min. Then, the eluted peptides from column were ionized and sprayed into the mass spectrometer, using a Nanospray Flex Ion Source ES071 (Thermo) under the following settings: spray voltage, 1.8 kV, capillary temperature, 250 °C. MS Raw data files were searched against the human protein sequence database or other species protein sequence database obtained from NCBI website using the Proteome Discoverer 1.4 software (Thermo, San Jose, CA) based on the SEQUEST and percolator algorithms. The false positive discovery rates (FDR) were set at 1 %. The resulting Proteome Discoverer Report contains all assembled proteins with peptides sequences and peptide spectrum match counts (PSM#) and TMT-tag based quantification ratio. TMT-tag based quantification was used for determining the relative abundance of proteins identified in each set of samples. The calculation and statistical analysis use Microsoft Excel functions. The heat map was generated using R. The annotation including pathways and processes was based on the Kyoto Encyclopedia of Genes and Genomes (KEGG) pathway database and UniProtKB protein database and the NCBI protein database. Samples were normalized to 353 proteins, which were used as control, due to no change between WT and *Wdr23KO* liver samples.

### HRS GeneWAS, population stratification, regression models and other covariates, and SNP evaluation

In brief, the US HRS[41, 42, 44] is a nationally representative, longitudinal sample of adults aged 50 years and older, who have been interviewed every 2 years, beginning in 1992. Because the HRS is nationally representative, including households across the country and the surveyed sample now includes over 36,000 participants, it is often used to calculate national prevalence rates for specific conditions for older adults, including physical and mental health outcomes, cognitive outcomes, as well as financial and social indicators. The sample for the current study is comprised of a subset of the HRS for which genetic data were collected, as described below. To reduce potential issues with population stratification, the GeneWAS in this study was limited to individuals of primarily European ancestry. The final sample was N = 3319, with the proportion of women at 58.5%.

#### HRS Participants

Data are from the *Health and Retirement Study* (HRS), a nationally representative sample of older Americans aged 50 and over[62, 63] in the contiguous United States. The present analysis was limited to participants who self-reported their race as white/Caucasian, verified by principal components analysis of ancestry markers, in order to assess effects of DCAF11 variation found in European ancestry groups. The analytical sample for the HRS included individuals who had available genetic data, at least one measure of hemoglobin A1C data, and relevant covariate data (N=9,326 to 9,333 per SNP based on sample and SNP quality).

#### DCAF11 Single Nucleotide Polymorphisms (SNPs)

For HRS, genotype data were accessed from the National Center for Biotechnology Information Genotypes and Phenotypes Database (dbGaP [64]). Genotyping was conducted on over 15,000 individuals using either the Illumina HumanOmni2.5-4v1 (2006 and 2008) and HumanOmni2.5-8v1 (2010) arrays and was performed by the NIH Center for Inherited Disease Research (CIDR). Standard quality control procedures were implemented by the University of Washington Genetic Coordinating Center [65]. Further detail is provided in HRS documentation [66]. The DCAF11 SNPs were filtered to include only those with a minor allele frequency of 5% or greater. SNPs were coded in order to assess additive effects of each additional allele (i.e., 0, 1, or 2 minor alleles) and were extracted using PLINK 1.9 [67, 68].

#### Hemoglobin A1C Biomarkers

The HRS collected biomarkers from blood spots, including glycosylated hemoglobin (HbA1c), which is an indicator of glycemic control over the past 2-3 months. HbA1c was available from blood spots on half of the sample in 2006, and the other half in 2008, with additional individuals captured in the 2010 or 2012 data collection waves. Detailed information on collection and assay are provided elsewhere [69, 70].

#### Covariates

In HRS, covariates included age at biomarker assessment, gender (0=female, 1=male), and four principal components to reduce such type 1 error due to differences in underlying population substructure[71, 72]. Detailed descriptions of the processes employed for running principal components analysis, including SNP selection, are provided by HRS[66], and follow methods outlined by Patterson and colleagues [73].

#### Statistical Analysis

All experiments were performed at least in triplicate. Data are presented as mean ± SEM. Data handling and statistical processing were performed using GraphPad Prism 8.0. Comparisons between two groups were done using unpaired Student’s t-test. Comparisons between more than two groups were done using the one-way ANOVA. Differences were considered significant at the p ≤ 0.05 level.

In HRS, multivariable linear regression models were run to test for the association between each of the five DCAF11 SNPs and hemoglobin A1C, adjusting for age, gender, and principal components using PLINK. With the number of SNPs and primary phenotypes in this study, strict Bonferroni correction would yield an adjusted multiple test-correction p-value threshold of 0.01 (for 5 SNP tests). However, a Bonferroni correction is too conservative for this type of gene-level assessment because of the correlations between SNPs within the gene [74, 75]. To address this, we calculate empirical p-value thresholds, through permutation procedures [74–77]. Permutation is a process whereby the correlations between SNPs and phenotypes are intentionally shuffled and then p-values calculated for the shuffled (null) data are compared to the non-shuffled data. This procedure is repeated multiple times in order to determine an empirical p-value [75, 77, 78], an empirically derived threshold at which a test result is less likely to achieve significance by chance alone. We performed 1,000 permutations using PLINK to derive the empirical p-value threshold of 0.0142 for HbA1c for determining statistically significant associations with DCAF11 SNPs.

## REFERENCES

1. Saeedi, P., et al., Global and regional diabetes prevalence estimates for 2019 and projections for 2030 and 2045: Results from the International Diabetes Federation Diabetes Atlas, 9(th) edition. Diabetes Res Clin Pract, 2019. 157: p. 107843.

2. Association, A.D., 2. Classification and Diagnosis of Diabetes:. Diabetes Care, 2021. 44(Suppl 1): p. S15–S33.

3. Najjar, S.M. and G. Perdomo, Hepatic Insulin Clearance: Mechanism and Physiology. Physiology (Bethesda), 2019. 34(3): p. 198–215.

4. Duckworth, W.C., R.G. Bennett, and F.G. Hamel, Insulin degradation: progress and potential. Endocr Rev, 1998. 19(5): p. 608–24.

5. Tokarz, V.L., P.E. MacDonald, and A. Klip, The cell biology of systemic insulin function. J Cell Biol, 2018. 217(7): p. 2273–2289.

6. BROH-KAHN, R.H. and I.A. MIRSKY, The inactivation of insulin by tissue extracts; the effect of fasting on the insulinase content of rat liver. Arch Biochem, 1949. 20(1): p. 10–4.

7. Corkey, B.E., Banting lecture 2011: hyperinsulinemia: cause or consequence? Diabetes, 2012. 61(1): p. 4–13.

8. Bojsen-Møller, K.N., et al., Hepatic Insulin Clearance in Regulation of Systemic Insulin Concentrations-Role of Carbohydrate and Energy Availability. Diabetes, 2018. 67(11): p. 2129–2136.

9. Pivovarova, O., et al., Hepatic insulin clearance is closely related to metabolic syndrome components. Diabetes Care, 2013. 36(11): p. 3779–85.

10. Shanik, M.H., et al., Insulin resistance and hyperinsulinemia: is hyperinsulinemia the cart or the horse? Diabetes Care, 2008. 31 Suppl 2: p. S262–8.

11. Asare-Bediako, I., et al., Variability of Directly Measured First-Pass Hepatic Insulin Extraction and Its Association With Insulin Sensitivity and Plasma Insulin. Diabetes, 2018. 67(8): p. 1495–1503.

12. Kotronen, A., et al., Increased liver fat, impaired insulin clearance, and hepatic and adipose tissue insulin resistance in type 2 diabetes. Gastroenterology, 2008. 135(1): p. 122–30.

13. Duckworth, W.C., Insulin degradation: mechanisms, products, and significance. Endocr Rev, 1988. 9(3): p. 319–45.

14. Yonezawa, K., et al., Insulin-degrading enzyme is capable of degrading receptor-bound insulin. Biochem Biophys Res Commun, 1988. 150(2): p. 605–14.

15. Villa-Pérez, P., et al., Liver-specific ablation of insulin-degrading enzyme causes hepatic insulin resistance and glucose intolerance, without affecting insulin clearance in mice. Metabolism, 2018. 88: p. 1–11.

16. Callis, J., The ubiquitination machinery of the ubiquitin system. Arabidopsis Book, 2014. 12: p. e0174.

17. Spatola, B.N., et al., Nuclear and cytoplasmic WDR-23 isoforms mediate differential effects on GEN-1 and SKN-1 substrates. Sci Rep, 2019. 9(1): p. 11783.

18. Zimmerman, E.S., B.A. Schulman, and N. Zheng, Structural assembly of cullin-RING ubiquitin ligase complexes. Curr Opin Struct Biol, 2010. 20(6): p. 714–21.

19. Lo, J.Y., B.N. Spatola, and S.P. Curran, WDR23 regulates NRF2 independently of KEAP1. PLoS Genet, 2017. 13(4): p. e1006762.

20. Brodersen, M.M., et al., CRL4(WDR23)-Mediated SLBP Ubiquitylation Ensures Histone Supply during DNA Replication. Mol Cell, 2016. 62(4): p. 627–35.

21. Titchenell, P.M., M.A. Lazar, and M.J. Birnbaum, Unraveling the Regulation of Hepatic Metabolism by Insulin. Trends Endocrinol Metab, 2017. 28(7): p. 497–505.

22. Shigiyama, F., et al., Mechanisms of sleep deprivation-induced hepatic steatosis and insulin resistance in mice. Am J Physiol Endocrinol Metab, 2018. 315(5): p. E848–e858.

23. Le, R., et al., Dcaf11 activates Zscan4-mediated alternative telomere lengthening in early embryos and embryonic stem cells. Cell Stem Cell, 2021. 28(4): p. 732–747.e9.

24. Xu, Z., et al., WDR-23 and SKN-1/Nrf2 Coordinate with the BLI-3 Dual Oxidase in Response to Iodide-Triggered Oxidative Stress. G3 (Bethesda), 2018. 8(11): p. 3515–3527.

25. Bazotte, R.B., L.G. Silva, and F.P. Schiavon, Insulin resistance in the liver: deficiency or excess of insulin? Cell Cycle, 2014. 13(16): p. 2494–500.

26. Leissring, M.A., Insulin-Degrading Enzyme: Paradoxes and Possibilities. Cells, 2021. 10(9).

27. Zhou, Z., M.J. Xu, and B. Gao, Hepatocytes: a key cell type for innate immunity. Cell Mol Immunol, 2016. 13(3): p. 301–15.

28. Lee, S. and H.H. Dong, FoxO integration of insulin signaling with glucose and lipid metabolism. J Endocrinol, 2017. 233(2): p. R67–r79.

29. Huang, X., et al., The PI3K/AKT pathway in obesity and type 2 diabetes. Int J Biol Sci, 2018. 14(11): p. 1483–1496.

30. Kay, A.M., C.L. Simpson, and J.A. Stewart, Jr., The Role of AGE/RAGE Signaling in Diabetes-Mediated Vascular Calcification. J Diabetes Res, 2016. 2016: p. 6809703.

31. Merino, B., et al., Hepatic insulin-degrading enzyme regulates glucose and insulin homeostasis in diet-induced obese mice. Metabolism, 2020. 113: p. 154352.

32. Mayer, A.L., et al., SLC2A8 (GLUT8) is a mammalian trehalose transporter required for trehalose-induced autophagy. Sci Rep, 2016. 6: p. 38586.

33. Guo, S., Molecular Basis of Insulin Resistance: The Role of IRS and Foxo1 in the Control of Diabetes Mellitus and Its Complications. Drug Discov Today Dis Mech, 2013. 10(1-2): p. e27–e33.

34. Probe Reports from the NIH Molecular Libraries Program. 2010.

35. Keenan, A.B., et al., ChEA3: transcription factor enrichment analysis by orthogonal omics integration. Nucleic Acids Res, 2019. 47(W1): p. W212–W224.

36. Tebay, L.E., et al., Mechanisms of activation of the transcription factor Nrf2 by redox stressors, nutrient cues, and energy status and the pathways through which it attenuates degenerative disease. Free Radic Biol Med, 2015. 88(Pt B): p. 108–146.

37. Uruno, A., et al., The Keap1-Nrf2 system prevents onset of diabetes mellitus. Mol Cell Biol, 2013. 33(15): p. 2996–3010.

38. Shaw, P. and A. Chattopadhyay, Nrf2-ARE signaling in cellular protection: Mechanism of action and the regulatory mechanisms. J Cell Physiol, 2020. 235(4): p. 3119–3130.

39. Tonelli, C., I.I.C. Chio, and D.A. Tuveson, Transcriptional Regulation by Nrf2. Antioxid Redox Signal, 2018. 29(17): p. 1727–1745.

40. Fisher, G.G. and L.H. Ryan, Overview of the Health and Retirement Study and Introduction to the Special Issue. Work Aging Retire, 2018. 4(1): p. 1–9.

41. Sonnega, A., et al., Cohort Profile: the Health and Retirement Study (HRS). Int J Epidemiol, 2014. 43(2): p. 576–85.

42. Juster, T. and R. Suzman, An Overview of the Health and Retirement Study, in Special Issue on the Health and Retirement Study: Data Quality and Early Results (1995). University of Wisconsin Press: The Journal of Human Resources. p. S7–S56.

43. Liu, Z., et al., Associations of genetics, behaviors, and life course circumstances with a novel aging and healthspan measure: Evidence from the Health and Retirement Study. PLoS Med, 2019. 16(6): p. e1002827.

44. Villa, O., et al., Genetic variation in. Elife, 2022. 11.

45. Lyons, T.J. and A. Basu, Biomarkers in diabetes: hemoglobin A1c, vascular and tissue markers. Transl Res, 2012. 159(4): p. 303–12.

46. Pivovarova, O., et al., Insulin-degrading enzyme: new therapeutic target for diabetes and Alzheimer’s disease? Ann Med, 2016. 48(8): p. 614–624.

47. Tang, W.J., Targeting Insulin-Degrading Enzyme to Treat Type 2 Diabetes Mellitus. Trends Endocrinol Metab, 2016. 27(1): p. 24–34.

48. Maianti, J.P., et al., Anti-diabetic activity of insulin-degrading enzyme inhibitors mediated by multiple hormones. Nature, 2014. 511(7507): p. 94–8.

49. Cotsapas, C., et al., Expression analysis of loci associated with type 2 diabetes in human tissues. Diabetologia, 2010. 53(11): p. 2334–9.

50. Hong, M.G., et al., Evidence that the gene encoding insulin degrading enzyme influences human lifespan. Hum Mol Genet, 2008. 17(15): p. 2370–8.

51. Titchenell, P.M., et al., Hepatic insulin signalling is dispensable for suppression of glucose output by insulin in vivo. Nat Commun, 2015. 6: p. 7078.

52. Wu, Y., et al., Novel Mechanism of Foxo1 Phosphorylation in Glucagon Signaling in Control of Glucose Homeostasis. Diabetes, 2018. 67(11): p. 2167–2182.

53. Yagishita, Y., et al., Nrf2 Improves Leptin and Insulin Resistance Provoked by Hypothalamic Oxidative Stress. Cell Rep, 2017. 18(8): p. 2030–2044.

54. Aleksunes, L.M., et al., Nuclear factor erythroid 2-related factor 2 deletion impairs glucose tolerance and exacerbates hyperglycemia in type 1 diabetic mice. The Journal of pharmacology and experimental therapeutics, 2010. 333(1): p. 140–151.

55. Vasileva, L.V., et al., Obesity and NRF2-mediated cytoprotection: Where is the missing link? Pharmacological Research, 2020. 156: p. 104760.

56. Li, L., et al., Glucose negatively affects Nrf2/SKN-1-mediated innate immunity in C. elegans. Aging, 2018. 10(11): p. 3089–3103.

57. Tang, L. and K.P. Choe, Characterization of skn-1/wdr-23 phenotypes in Caenorhabditis elegans; pleiotrophy, aging, glutathione, and interactions with other longevity pathways. Mech Ageing Dev, 2015. 149: p. 88–98.

58. Choe, K.P., A.J. Przybysz, and K. Strange, The WD40 repeat protein WDR-23 functions with the CUL4/DDB1 ubiquitin ligase to regulate nuclear abundance and activity of SKN-1 in Caenorhabditis elegans. Molecular and cellular biology, 2009. 29(10): p. 2704–15.

59. Ryder, E., et al., Molecular characterization of mutant mouse strains generated from the EUCOMM/KOMP-CSD ES cell resource. Mamm Genome, 2013. 24(7-8): p. 286–94.

60. White, J.K., et al., Genome-wide generation and systematic phenotyping of knockout mice reveals new roles for many genes. Cell, 2013. 154(2): p. 452–64.

61. Benedé-Ubieto, R., et al., Guidelines and Considerations for Metabolic Tolerance Tests in Mice. Diabetes Metab Syndr Obes, 2020. 13: p. 439–450.

62. HRS, Sample Sizes and Response Rates. 2011, University of Michigan: Ann Arbor, MI. p. 1–13. http://hrsonline.isr.umich.edu/sitedocs/sampleresponse.pdf (Accessed October 12, 2016).

63. Juster, F.T. and R. Suzman, An overview of the Health and Retirement Study. Journal of Human Resources, 1995: p. S7–S56.

64. dbGaP, Health and Retirement Study. 2012, National Center for Biotechnology Information: Bethesda, MD.

65. Laurie, C.C., et al., Quality control and quality assurance in genotypic data for genome-wide association studies. Genetic epidemiology, 2010. 34(6): p. 591–602.

66. HRS, Quality control report for genotypic data. 2012, University of Washington: St. Louis, MO. p. 1–44. http://hrsonline.isr.umich.edu/sitedocs/genetics/HRS_QC_REPORT_MAR2012.pdf (Accessed March 15, 2015).

67. Chang, C.C., et al., Second-generation PLINK: rising to the challenge of larger and richer datasets. arXiv preprint arXiv:1410.4803, 2014.

68. Purcell, S., et al., PLINK: a tool set for whole-genome association and population-based linkage analyses. American journal of human genetics, 2007. 81(3): p. 559–75.

69. Crimmins, E., et al., Documentation of biomarkers in the 2006 and 2008 Health and Retirement Study. Ann Arbor, MI: Survey Research Center University of Michigan, 2013.

70. Crimmins, E., et al., Validation of blood-based assays using dried blood spots for use in large population studies. Biodemography and social biology, 2014. 60(1): p. 38–48.

71. Tian, C., P.K. Gregersen, and M.F. Seldin, Accounting for ancestry: population substructure and genome-wide association studies. Human molecular genetics, 2008. 17(R2): p. R143–R150.

72. Price, A.L., et al., Principal components analysis corrects for stratification in genome-wide association studies. Nature genetics, 2006. 38(8): p. 904–909.

73. Patterson, N., A.L. Price, and D. Reich, Population structure and eigenanalysis. PLoS genetics, 2006. 2(12): p. e190.

74. Han, B., H.M. Kang, and E. Eskin, Rapid and accurate multiple testing correction and power estimation for millions of correlated markers. PLoS Genet, 2009. 5(4): p. e1000456.

75. Sham, P.C. and S.M. Purcell, Statistical power and significance testing in large-scale genetic studies. Nat Rev Genet, 2014. 15(5): p. 335–46.

76. Dudbridge, F. and A. Gusnanto, Estimation of significance thresholds for genomewide association scans. Genet Epidemiol, 2008. 32(3): p. 227–34.

77. Pahl, R. and H. Schafer, PERMORY: an LD-exploiting permutation test algorithm for powerful genome-wide association testing. Bioinformatics, 2010. 26(17): p. 2093–100.

78. North, B.V., D. Curtis, and P.C. Sham, A note on the calculation of empirical P values from Monte Carlo procedures. Am J Hum Genet, 2002. 71(2): p. 439–41.

