## Supplemental Material for "Hepatic WDR23 proteostasis mediates insulin clearance by regulating insulin-degrading enzyme"

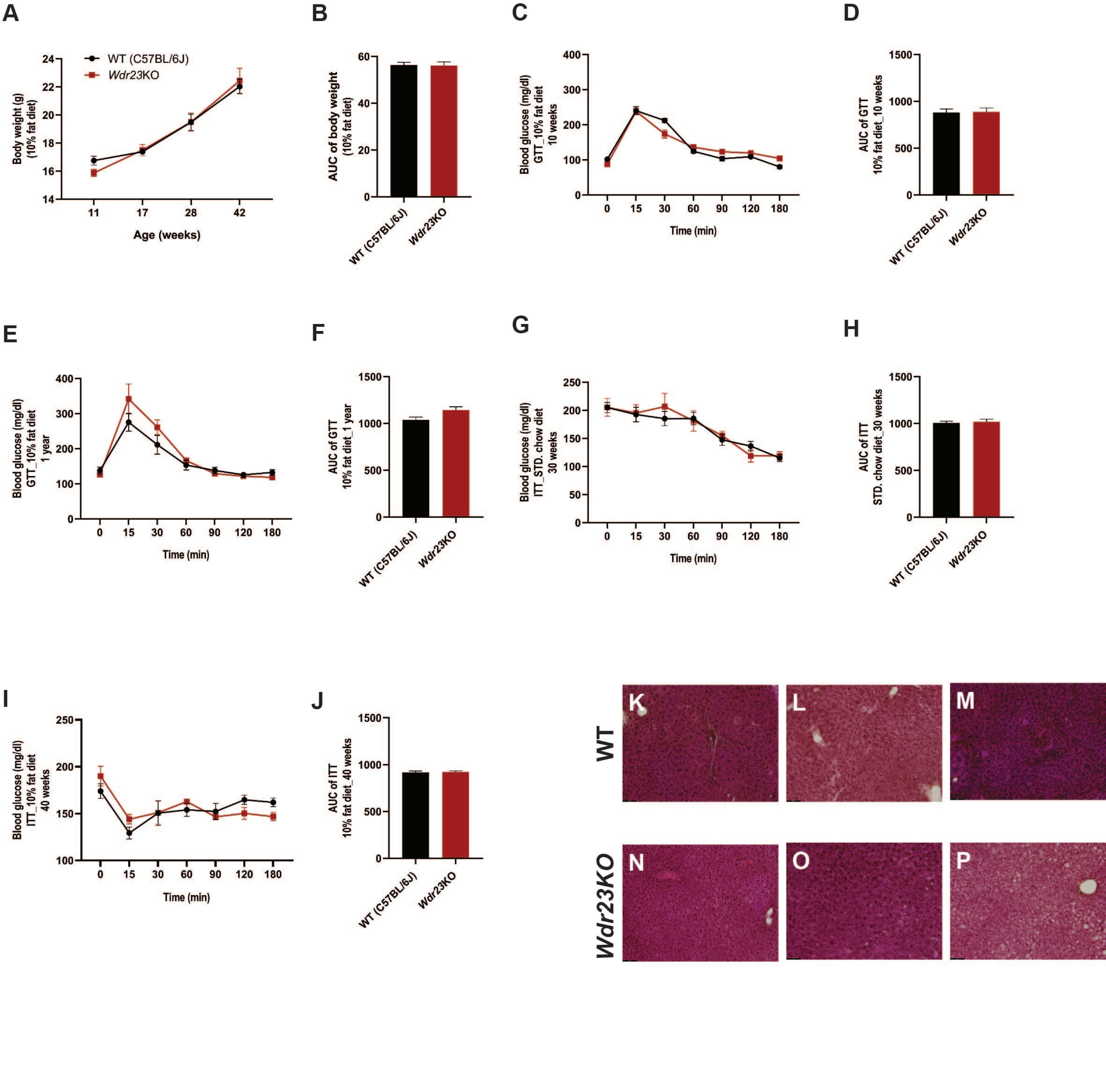

Duangjan Supplemental Figure 1

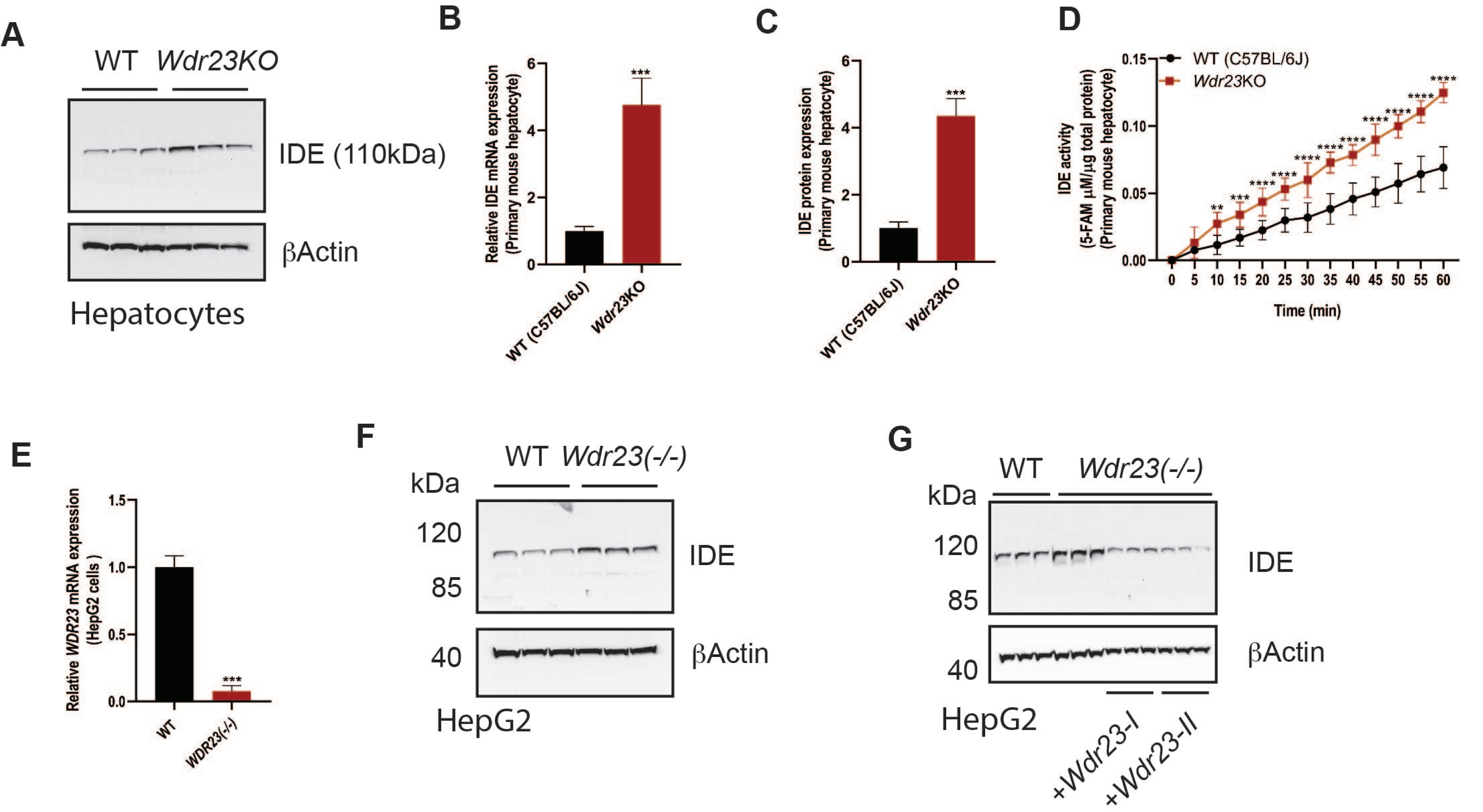

Duangjan Supplemental Figure 2

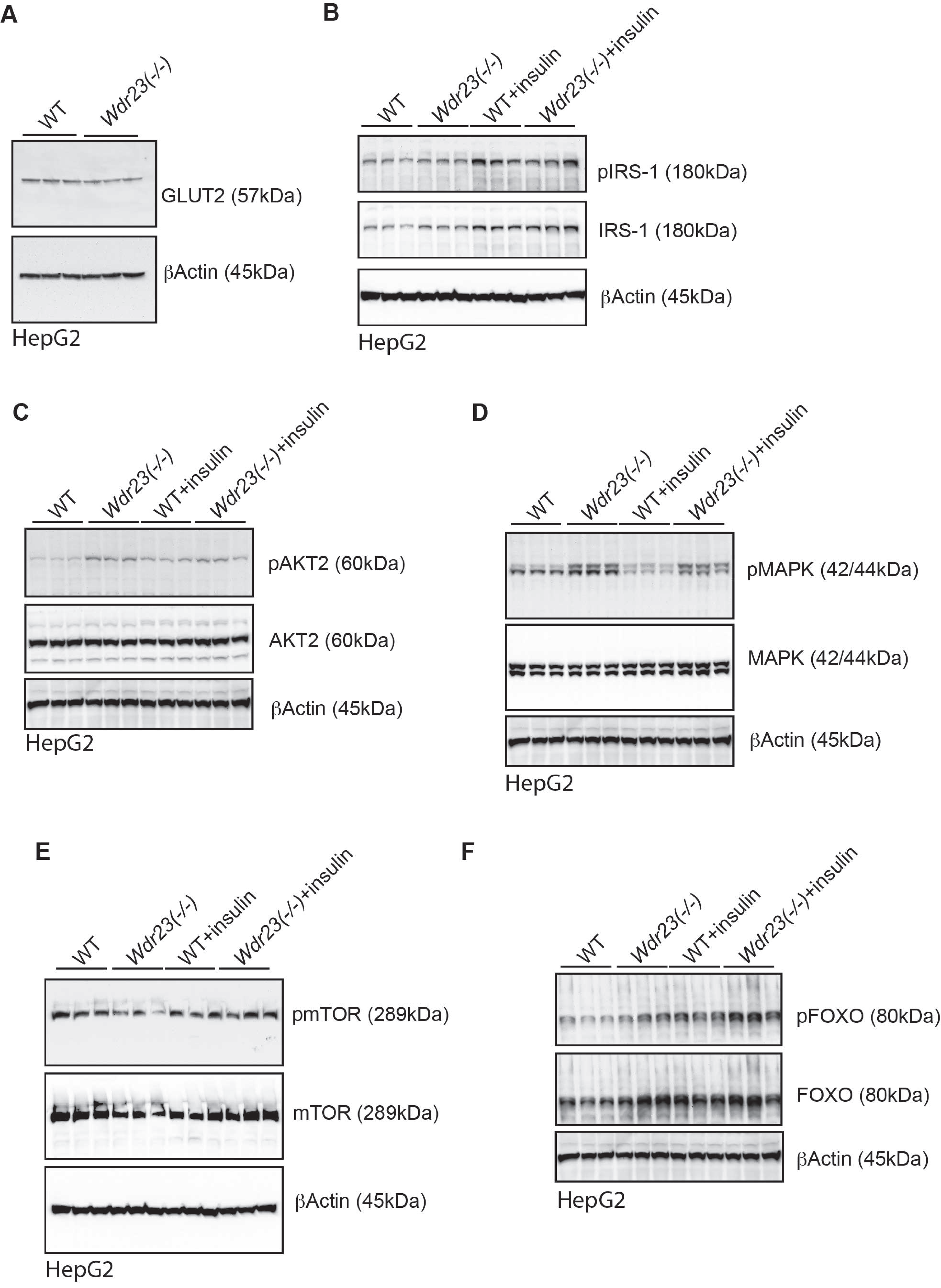

Duangjan Supplemental Figure 3

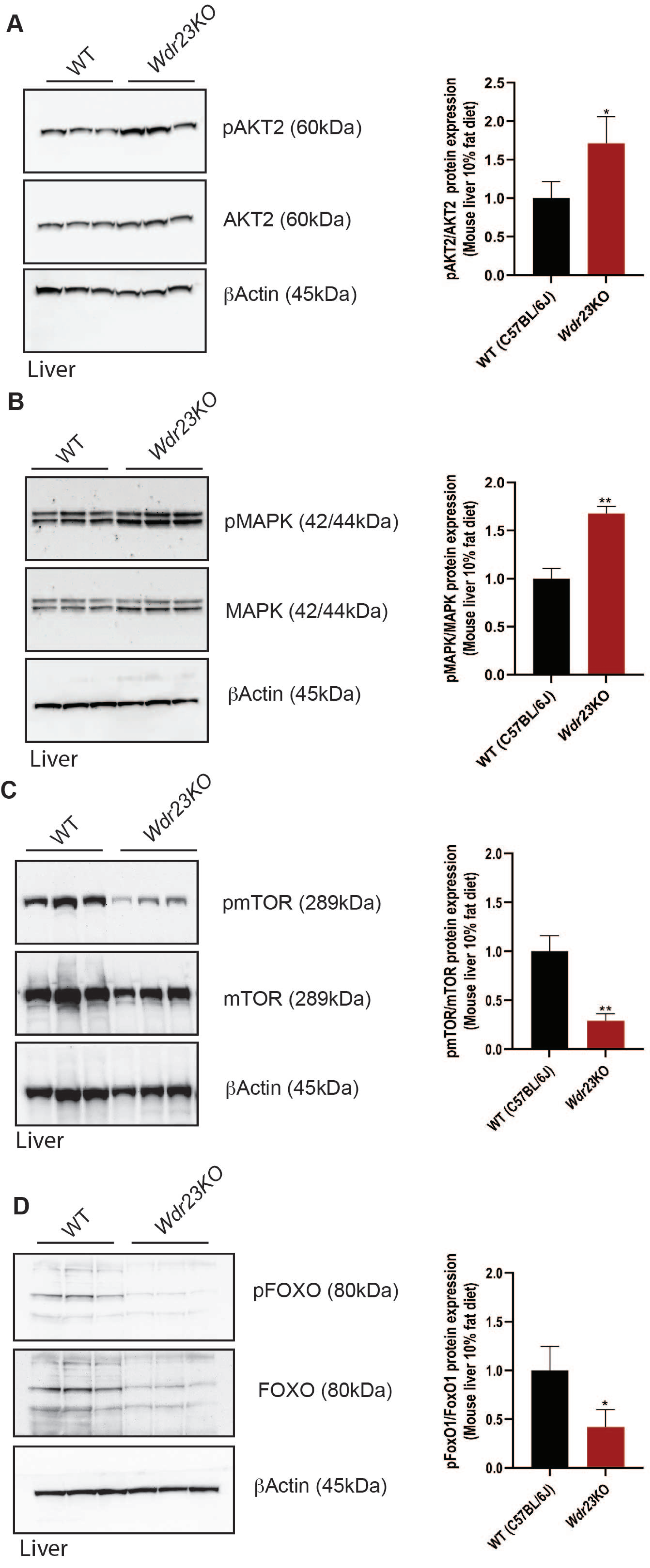

Duangjan Supplemental Figure 4

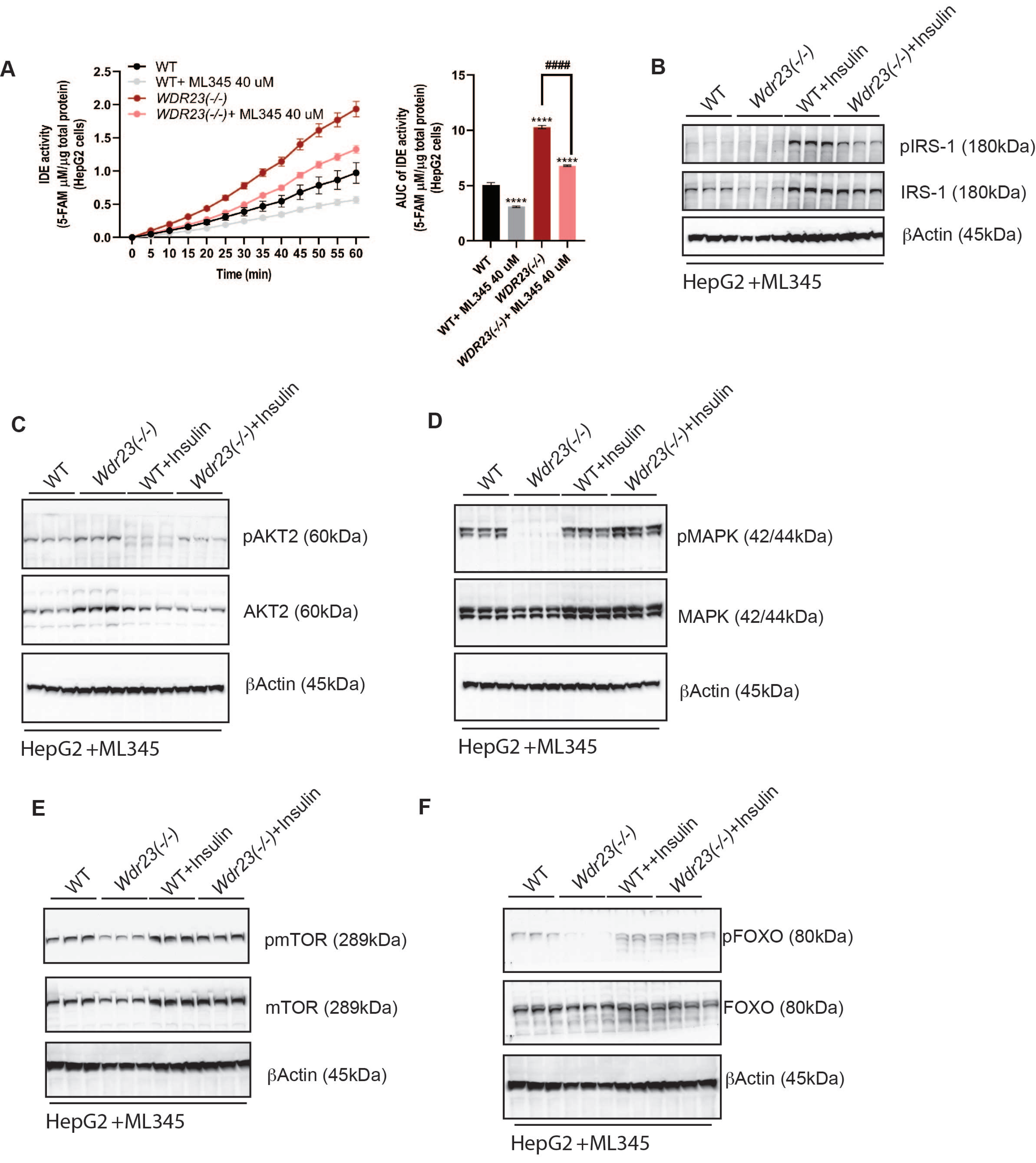

Duangjan Supplemental Figure 5

A

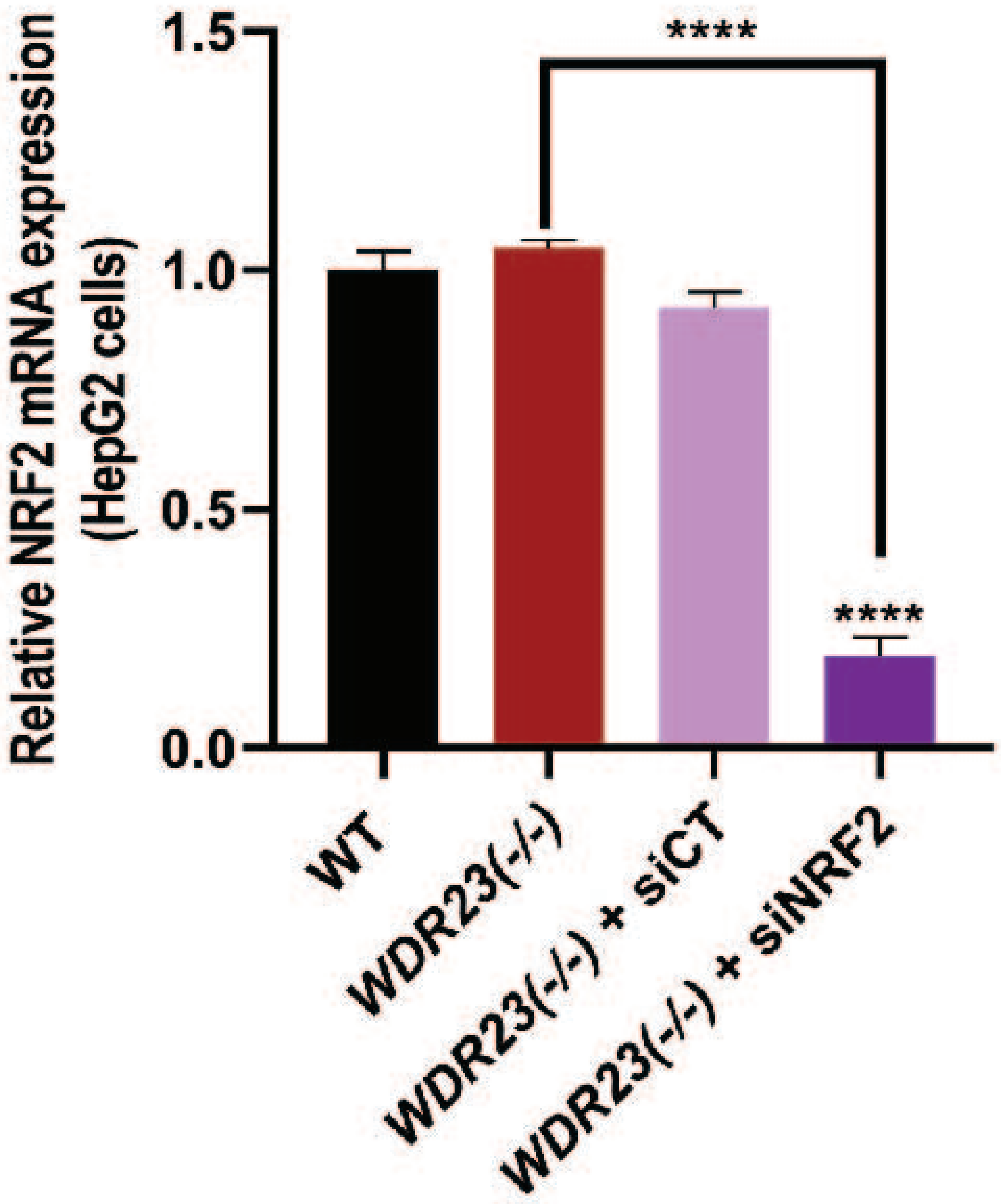

B

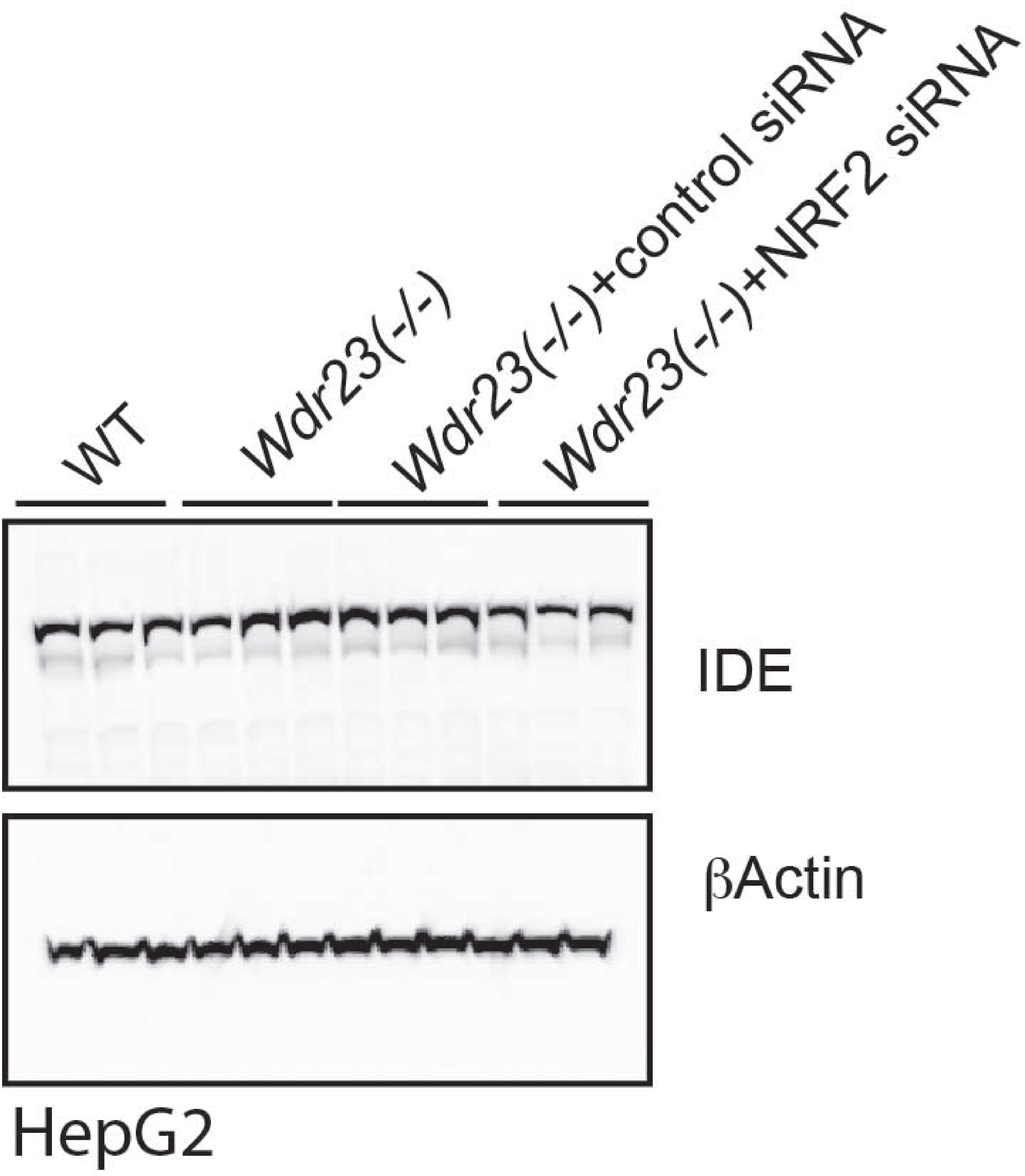

Duangjan Supplemental Figure 6

**Table S1. The GO functional enrichment analysis of DEGs in *Wdr23*KO mice liver samples compare to the WT control with the threshold of *P*≤0.05**

| Term | Gene ID | Chr | Name | Description | Log2 Fold Change | *P*-value | *P*-adj |
|---|---|---|---|---|---|---|---|
| **Carbohydrate metabolic process Up-regulated** | | | | | | | |
| | ENSMUSG00000041237 | 3 | Pklr | pyruvate kinase liver and red blood cell | 1.842783 | 3.55E-10 | 5.68E-07 |
| | ENSMUSG00000004815 | 5 | Dgkq | diacylglycerol kinase | 2.009794 | 1.57E-08 | 1.33E-05 |
| | ENSMUSG00000041798 | 11 | Gck | glucokinase | 1.839161 | 5.97E-06 | 0.00141 |
| | ENSMUSG00000025815 | 2 | Dhtkd1 | dehydrogenase E1 and transketolase domain containing 1 | 1.114049 | 7.15E-05 | 0.008823 |
| | ENSMUSG00000029802 | 6 | Abcg2 | ATP binding cassette subfamily G member 2 (Junior blood group) | 1.101674 | 0.000264 | 0.021585 |
| | ENSMUSG00000034793 | 11 | G6pc3 | glucose 6 phosphatase, catalytic, 3 | 1.298063 | 0.000421 | 0.030001 |
| | ENSMUSG00000025236 | 9 | Adpgk | ADP-dependent glucokinase | 1.300949 | 0.00093 | 0.048996 |
| | ENSMUSG00000041237 | 3 | Pklr | pyruvate kinase liver and red blood cell | 1.842783 | 3.55E-10 | 5.68E-07 |
| | ENSMUSG00000025815 | 2 | Dhtkd1 | dehydrogenase E1 and transketolase domain containing 1 | 1.114049 | 7.15E-05 | 0.008823 |
| | ENSMUSG00000032310 | 9 | Cyp1a2 | cytochrome P450, family 1, subfamily a, polypeptide 2 | 1.524902 | 0.000215 | 0.018769 |
| | ENSMUSG00000025153 | 11 | Fasn | fatty acid synthase | 1.817651 | 0.000268 | 0.021806 |
| | ENSMUSG00000030972 | 7 | Acsm5 | acyl-CoA synthetase medium-chain family member 5 | 1.005508 | 0.000429 | 0.030242 |
| | ENSMUSG00000025236 | 9 | Adpgk | ADP-dependent glucokinase | 1.300949 | 0.00093 | 0.048996 |

| | | | | | | | |
|---|---|---|---|---|---|---|---|
| **Carbohydrate metabolic process**<br>**Down-regulated** | | | | | | | |
| | ENSMUSG00000000628 | 6 | Hk2 | hexokinase 2 | -3.22647 | 5.12E-06 | 0.001263 |
| | ENSMUSG00000024042 | 17 | Sik1 | salt inducible kinase 1 | -1.72827 | 1.38E-05 | 0.002671 |
| | ENSMUSG00000025190 | 19 | Got1 | glutamic-oxaloacetic transaminase 1 | -1.30683 | 7.23E-05 | 0.008859 |
| | ENSMUSG00000060402 | 7 | Chst8 | carbohydrate (N-acetylgalactosamine 4-0) sulfotransferase 8 | -7.0051 | 0.000148 | 0.014403 |
| | ENSMUSG00000024029 | 17 | Tff3 | trefoil factor 3, intestinal | -7.60111 | 0.000173 | 0.015887 |
| | ENSMUSG00000038155 | 19 | Gstp2 | glutathione S-transferase, pi 2 | -3.93441 | 3.93E-11 | 7.12E-08 |
| | ENSMUSG00000030895 | 7 | Hpx | hemopexin | -1.05529 | 7.74E-06 | 0.001801 |
| | ENSMUSG00000018339 | 11 | Gpx3 | glutathione peroxidase 3 | -1.63836 | 0.000341 | 0.025717 |
| | ENSMUSG00000031584 | 8 | Gsr | glutathione reductase | -1.10922 | 0.000844 | 0.046274 |
| **Fatty acid metabolic process**<br>**Up-regulated** | | | | | | | |
| | ENSMUSG00000020538 | 11 | Srebf1 | sterol regulatory element binding transcription factor 1 | 2.246853 | 2.76E-07 | 0.000145 |
| | ENSMUSG00000038754 | 19 | Elovl3 | elongation of very long chain fatty acids (FEN1/Elo2, SUR4/Elo3, yeast)-like 3 | 1.941961 | 6.96E-07 | 0.000298 |
| | ENSMUSG00000054422 | 6 | Fabp1 | fatty acid binding protein 1, liver | 1.256687 | 0.000107 | 0.011728 |
| | ENSMUSG00000025153 | 11 | Fasn | fatty acid synthase | 1.817651 | 0.000268 | 0.021806 |
| | ENSMUSG00000030972 | 7 | Acsm5 | acyl-CoA synthetase medium-chain family member 5 | 1.005508 | 0.000429 | 0.030242 |
| | ENSMUSG00000010651 | 9 | Acaa1b | acetyl-Coenzyme A acyltransferase 1B | 0.952087 | 0.000938 | 0.049223 |
| | ENSMUSG00000041798 | 11 | Gck | glucokinase | 1.839161 | 5.97E-06 | 0.00141 |
| | ENSMUSG00000040505 | 17 | Abcg5 | ATP binding | 1.47019 | 0.00028 | 0.022564 |

| | | | | | | | |
|---|---|---|---|---|---|---|---|
| | | | | cassette subfamily G member 5 | | | |
| | ENSMUSG00000047822 | 9 | Angptl8 | angiopoietin-like 8 | 1.571686 | 0.000583 | 0.036294 |
| | ENSMUSG00000005677 | 1 | Nr1i3 | nuclear receptor subfamily 1, group I, member 3 | 2.030792 | 0.000657 | 0.03918 |
| **Fatty acid metabolic process Down-regulated** | | | | | | | |
| | ENSMUSG00000039202 | 7 | Abhd2 | abhydrolase domain containing 2 | -2.58157 | 1.2E-05 | 0.002434 |
| | ENSMUSG00000015568 | 8 | Lpl | lipoprotein lipase | -2.73116 | 1.56E-05 | 0.002854 |
| | ENSMUSG00000074254 | 7 | Cyp2a4 | cytochrome P450, family 2, subfamily a, polypeptide 4 | -7.31411 | 0.000103 | 0.0116 |
| | ENSMUSG00000028341 | 4 | Nr4a3 | nuclear receptor subfamily 4, group A, member 3 | -7.15208 | 0.000104 | 0.01162 |
| | ENSMUSG00000031278 | X | Acsl4 | acyl-CoA synthetase long-chain family member 4 | -1.29954 | 0.000762 | 0.042973 |
| | ENSMUSG00000046402 | 9 | Rbp1 | retinol binding protein 1, cellular | -1.94839 | 2.09E-08 | 1.55E-05 |
| | ENSMUSG00000015568 | 8 | Lpl | lipoprotein lipase | -2.73116 | 1.56E-05 | 0.002854 |
| | ENSMUSG00000025190 | 19 | Got1 | glutamic-oxaloacetic transaminase 1, soluble | -1.30683 | 7.23E-05 | 0.008859 |
| | ENSMUSG00000021242 | 12 | Npc2 | NPC intracellular cholesterol transporter 2 | -1.26822 | 0.000168 | 0.015593 |

**Table S2. The KEGG pathway enrichment analysis of DEGs in *Wdr23*KO mice liver samples compare to the WT control with the threshold of *P*≤0.05**

| Term | Gene ID | Chr | Name | Description | Log2 Fold Change | *P*-value | *P*-adj |
|---|---|---|---|---|---|---|---|
| **Insulin signaling pathway Up-regulated** | | | | | | | |
| | ENSMUSG00000041237 | 3 | Pklr | pyruvate kinase liver and red blood cell | 1.842783 | 3.55E-10 | 5.68E-07 |
| | ENSMUSG00000020538 | 11 | Srebf1 | sterol regulatory element binding transcription factor 1 | 2.246853 | 2.76E-07 | 0.000145 |
| | ENSMUSG00000041798 | 11 | Gck | glucokinase | 1.839161 | 5.97E-06 | 0.00141 |
| | ENSMUSG00000025153 | 11 | Fasn | fatty acid synthase | 1.817651 | 0.000268 | 0.021806 |
| | ENSMUSG00000034793 | 11 | G6pc3 | glucose 6 phosphatase, catalytic, 3 | 1.298063 | 0.000421 | 0.030001 |
| | ENSMUSG00000020538 | 11 | Srebf1 | sterol regulatory element binding transcription factor 1 | 2.246853 | 2.76E-07 | 0.000145 |
| | ENSMUSG00000022383 | 15 | Ppara | peroxisome proliferator activated receptor alpha | 1.19511 | 0.001098 | 0.054689 |
| | ENSMUSG00000028978 | 5 | Nos3 | nitric oxide synthase 3, endothelial cell | 1.637867 | 0.00291 | 0.09853 |
| **MAPK signaling pathway Up-regulated** | | | | | | | |
| | ENSMUSG00000054252 | 5 | Fgfr3 | fibroblast growth factor receptor 3 | 1.303898 | 0.000326 | 0.025083 |
| **FoxO signaling pathway Up-regulated** | | | | | | | |
| | ENSMUSG00000034793 | 11 | G6pc3 | glucose 6 phosphatase, catalytic, 3 | 1.298063 | 0.000421 | 0.030001 |
| **Glycolysis / Gluconeogenesis Up-regulated** | | | | | | | |
| | ENSMUSG00000041237 | 3 | Pklr | pyruvate kinase liver and red blood cell | 1.842783 | 3.55E-10 | 5.68E-07 |
| | ENSMUSG00000041798 | 11 | Gck | glucokinase | 1.839161 | 5.97E-06 | 0.00141 |
| | ENSMUSG00000034793 | 11 | G6pc3 | glucose 6 phosphatase, catalytic, 3 | 1.298063 | 0.000421 | 0.030001 |

| | | | | | | | |
|---|---|---|---|---|---|---|---|
| | ENSMUSG00000025236 | 9 | Adpgk | ADP-dependent glucokinase | 1.300949 | 0.00093 | 0.048996 |
| **Pyruvate metabolism<br>Up-regulated** | | | | | | | |
| | ENSMUSG00000041237 | 3 | Pklr | pyruvate kinase liver and red blood cell | 1.842783 | 3.55E-10 | 5.68E-07 |
| **PPAR signaling pathway<br>Up-regulated** | | | | | | | |
| | ENSMUSG00000054422 | 6 | Fabp1 | fatty acid binding protein 1, liver | 1.256687 | 0.000107 | 0.011728 |
| | ENSMUSG00000010651 | 9 | Acaa1b | acetyl-Coenzyme A acyltransferase 1B | 0.952087 | 0.000938 | 0.049223 |
| **AGE-RAGE signaling pathway<br>Down-regulated** | | | | | | | |
| | ENSMUSG00000027962 | 3 | Vcam1 | vascular cell adhesion molecule 1 | -1.33177 | 2.8E-05 | 0.004321 |
| **Glutathione metabolism<br>Down-regulated** | | | | | | | |
| | ENSMUSG00000038155 | 19 | Gstp2 | glutathione S-transferase, pi 2 | -3.93441 | 3.93E-11 | 7.12E-08 |
| | ENSMUSG00000004038 | 3 | Gstm3 | glutathione S-transferase, mu 3 | -3.42511 | 0.000332 | 0.025253 |
| | ENSMUSG00000018339 | 11 | Gpx3 | glutathione peroxidase 3 | -1.63836 | 0.000341 | 0.025717 |
| | ENSMUSG00000028597 | 4 | Gpx7 | glutathione peroxidase 7 | -3.05796 | 0.000596 | 0.036628 |
| | ENSMUSG00000031584 | 8 | Gsr | glutathione reductase | -1.10922 | 0.000844 | 0.046274 |

**Table S3. Differential expression of targeted proteins in *Wdr23*KO mice liver samples compare to the WT control with the threshold of *P*≤0.05**

| Term | Name | Description | Log2 Fold Change | *P*-value | *P*-adj |
|---|---|---|---|---|---|
| **Carbohydrate metabolism Up-regulated** | | | | | |
| | SNX4 | Sorting nexin-4 | 0.083229 | 1.059386 | 0.00123 |
| | GTR8/SLC2A8 | Glucose transporter type 8 | 0.123433 | 1.089324 | 0.047594 |
| | RET4/RBP4 | Retinol-binding protein 4 | 0.164573 | 1.120834 | 0.015476 |
| | VPS39 | Vam6/Vps39-like protein | 0.239837 | 1.180859 | 0.03333 |
| | IDE | Insulin-degrading enzyme | 0.49103 | 1.405448 | 0.003979 |
| | PLIN2 | Perilipin-2 | 0.595647 | 1.51115 | 0.044223 |
| | CBPB2 | Carboxypeptidase B2 | 0.122307 | 1.088474 | 0.044443 |
| | GALE | UDP-glucose 4-epimerase | 0.202877 | 1.150991 | 0.002718 |
| **Carbohydrate metabolism Down-regulated** | | | | | |
| | HRG | Haeme-responsive gene (HRG)-1 | -0.6263 | 0.647837 | 0.020691 |
| | NAGTC | Sodium-dependent glucose transporter 1C | -0.23073 | 0.852203 | 0.016698 |
| | C2C2L | Transmembrane protein 24 | -0.14151 | 0.906567 | 0.03487 |
| | ACVR1/TGF | TGF-B superfamily receptor type I | -0.13201 | 0.91256 | 0.027382 |
| | LMAN1 | Lectin mannose-binding 1 | -0.12298 | 0.918291 | 0.013709 |
| | RAB10 | Ras-related protein Rab-10 | -0.08957 | 0.939802 | 0.016464 |
| | XDH | Xanthine dehydrogenase | -0.11324 | 0.92451 | 0.004215 |
| **Insulin signaling Up-regulated** | | | | | |
| | PEDF | Serpin F1 | 0.078545 | 1.055952 | 0.021617 |
| | SORCN | Sorcin | 0.174471 | 1.128551 | 0.023325 |
| **Insulin signaling Down-regulated** | | | | | |
| | LMAN2 | Lectin mannose-binding 2 | -0.19273 | 0.874951 | 0.036281 |
| | BAIP2/IRSp53 | Insulin receptor substrate protein of 53 kDa | -0.13084 | 0.913297 | 0.001019 |
| | OGT1 | O-GlcNAc transferase subunit p110 | -0.09375 | 0.937081 | 0.012789 |
| **PI3K/AKT/mTOR signaling Up-regulated** | | | | | |
| | MTOR | Serine/threonine-protein kinase mTOR | 0.071131 | 1.05054 | 0.010672 |
| | LTOR1 | Late endosomal/lysosomal adaptor and MAPK and MTOR activator 1 | 0.08185 | 1.058374 | 0.049572 |
| | GRP75 | Glucose-regulated protein 75 (GRP75), Heat shock protein 70 (HSP70) | 0.091203 | 1.065258 | 0.021406 |
| | 1433Z | Protein kinase C inhibitor protein 1 | 0.135461 | 1.098444 | 0.015504 |
| | ASNS | Glutamine-dependent asparagine synthetase | 1.777318 | 3.427883 | 0.022428 |
| **PI3K/AKT/mTOR signaling Down-regulated** | | | | | |
| | SCLY | Selenocysteine lyase | -0.21566 | 0.861155 | 0.020388 |
| | NRN1 | Neuritin | -0.40308 | 0.756242 | 0.004372 |
| | AKTS1/PRAS | Proline-rich AKT1 substrate 1 | -0.12297 | 0.918296 | 0.015158 |
| **FoxO signaling Up-regulated** | | | | | |
| | FCOR | Foxo1-corepressor | 0.421892 | 1.339683 | 0.000777 |

| | | | | | |
|---|---|---|---|---|---|
| **FoxO signaling Down-regulated** | | | | | |
| | HMGA1 | High mobility group protein HMG-I | -0.33595 | 0.792261 | 0.003751 |
| **MAPK signaling Up-regulated** | | | | | |
| | MK09 | MAPK9 | 0.110373 | 1.079507 | 0.000505 |
| | ECSIT | SITPEC, Evolutionarily conserved signaling intermediate in Toll pathway | 0.115942 | 1.083682 | 0.025137 |
| **MAPK signaling Down-regulated** | | | | | |
| | M3K3 | MAPK/ERK kinase kinase 3 | -0.6179 | 0.651617 | 0.032129 |
| | PAXI | Paxillin | -0.13104 | 0.91317 | 0.002378 |
| | GAB1 | GRB2-associated-binding protein 1 | -0.07947 | 0.946408 | 0.006037 |
| | SASH1 | SAM and SH3 domain-containing protein 1 | -0.11865 | 0.921048 | 0.027416 |
| **AMPK signaling Down-regulated** | | | | | |
| | CRBN | Cereblon | -0.93098 | 0.524503 | 0.010892 |
| | PARP1 | Poly [ADP-ribose] polymerase 1 | -0.08163 | 0.94499 | 0.029268 |
| | TBK1 | TANK-binding kinase 1 | -0.47323 | 0.720352 | 0.037901 |
| **Lipid metabolism Up-regulated** | | | | | |
| | ASAH1 | Acid ceramidase | 0.246443 | 1.186279 | 0.04491 |
| | THIKB | Beta-ketothiolase B | 0.349508 | 1.274126 | 0.04113 |
| **Lipid metabolism Down-regulated** | | | | | |
| | ELOV1 | Elongation of very long chain fatty acids protein 1 | -0.441 | 0.736622 | 0.010616 |
| | AAKG2 | AMP-activated protein kinase subunit gamma-2 | -0.43151 | 0.741486 | 0.045831 |
| | OSBL2 | Oxysterol-binding protein-related protein 2 | -0.32967 | 0.795717 | 0.027604 |
| **Oxidation-reduction Up-regulation** | | | | | |
| | NDUF5 | NADH dehydrogenase [ubiquinone] 1 alpha subcomplex assembly factor 5 | 0.28191 | 1.215804 | 0.027516 |
| | NNTM | Nicotinamide nucleotide transhydrogenase | 2.126628 | 4.366955 | 5.49E-06 |
| | TMLH | Trimethyllysine dioxygenase | 0.320616 | 1.248864 | 0.014325 |
| **Oxidation-reduction Down-regulation** | | | | | |
| | COX19 | Cytochrome c oxidase assembly protein COX19 | -0.33592 | 0.792279 | 0.008761 |
| | AOFA | Amine oxidase [flavin-containing] A | -0.33583 | 0.79233 | 0.047069 |

| Table S4 | | | | |
|---|---|---|---|---|
| **Protein** | **Ratio (KO/WT)** | **p-value** | **Description** | **Role** |
| YWHAZ | 1.098443735 | 0.015503955 | 14-3-3 protein zeta/delta | Glucose Metabolism[1] |
| FCOR | 1.339683216 | 0.00077717 | Foxo1-corepressor and cellular glucose homeostasis | Transcriptional regulation [2] |
| AAKG2 | 0.741485544 | 0.045830957 | AMP/ATP-binding subunit of AMP-activated protein kinase (AMPK), an energy sensor protein kinase that plays a key role in regulating cellular energy metabolism. | Glucose metabolism[3] |
| IMPA2 | 1.141395562 | 0.000643951 | inositol monophosphatase 2 | Metabolic signaling[4] |
| PGP | 1.095449769 | 0.039508474 | Phosphoglycolate phosphatase | Healthy aging [5] |
| MTOR | 1.05054021 | 0.010672401 | mechanistic target of rapamycin kinase | Metabolic signaling[6] |
| CBP2 | 1.088474324 | 0.044443196 | Prohormone processing enzyme; carboxypeptidase | Metabolic homeostasis[7] |
| SLC2A8 | 1.089323636 | 0.047593704 | solute carrier family 2, (facilitated glucose transporter), member 8 | Carbohydrate transporter[8] |
| GALE | 1.150991471 | 0.00271846 | UDP-glucose 4-epimerase | Carbohydrate metabolism[9] |
| ASNS | 3.42788289 | 0.022427988 | asparagine synthetase | Response to glucose restriction |
| IDE | 1.405447674 | 0.003978549 | Insulin degrading enzyme | Insulin degradation[10,11] |
